# A general and extensible algorithmic framework to biological sequence alignment across scales and applications

**DOI:** 10.64898/2026.01.28.702355

**Authors:** Hao Xuan, Hongyang Sun, Xiangtao Liu, Hanyuan Zhang, Jun Zhang, Cuncong Zhong

## Abstract

Sequence alignment underpins nearly every facet of modern genomics, from genetic testing and cancer profiling to functional genome annotation. Yet, despite decades of algorithmic innovation, most existing aligners remain narrowly optimized for specific tasks, fragmenting analytical workflows and limiting reproducibility. Here we introduce the Versatile Alignment Toolkit (VAT), a unified algorithmic framework that generalizes existing seeding and genome-indexing strategies within a single, transparent architecture. VAT employs a novel multi-view indexing scheme that integrates multiple seeding strategies and supports run-time seed-length parameterization without reindexing. A radix clustering algorithm based on hardware-efficient in-register bitonic sort accelerates multi-view table construction, ensuring scalability across large datasets. VAT delivers consistently high performance across diverse alignment tasks, including short- and long-read mapping, homology search, and whole-genome alignment, while maintaining algorithmic simplicity and adaptability. By bridging previously isolated alignment paradigms, VAT substantially reduces workflow complexity, enhances computational efficiency, and establishes an extensible foundation for future sequencing technologies. We anticipate this unification will set a new standard for flexible and reproducible sequence alignment in biomedical research and clinical genomics.

## 1. Introduction

Sequence alignment, the quantitative assessment of similarity between biological sequences, forms the foundation of modern computational biology. Initially developed to infer gene and protein functions and to reconstruct evolutionary relationships, forming the foundation of comparative and functional genomics, sequence alignment now further underpins the analysis of high-throughput sequencing (HTS) data across a wide range of applications such as genomics, transcriptomics, and epigenomics^1-7^. The rapid expansion of sequencing capacity has generated biological data at an unprecedented scale, transforming alignment from a comparative tool into a critical gateway for biological discovery and clinical translation^8-10^. As sequence-derived information increasingly guides precision medicine, efficient and accurate sequence alignment has become indispensable to both basic research and biomedical applications^11-16^.

Sequence alignment was originally formulated as an edit-distance minimization problem and solved by dynamic programming algorithms such as Needleman–Wunsch and Smith–Waterman^17-20^. However, their quadratic time complexity rendered them impractical for large-scale datasets, prompting the development of heuristic, seed-based approaches that restrict alignment to regions of high local similarity^21-25^. Although this filtering concept is broadly applicable, real-world sequence data vary widely in alphabet (nucleotide and peptide), length (from short reads to whole genomes), structure (contiguous or spliced), expected similarity (from distant homologs to nearly identical reads), and scale^26-28^. To cope with this diversity, modern aligners specialize through tailored seeding (e.g., exact match, spaced seed^29^, minimizer^30-34^, or reduced alphabet^35^) and indexing strategies (e.g., hashing^36,37^, suffix arrays^38,39^, Burrows–Wheeler transform^40-42^, or double indexing^43,44^). While these task-specific optimizations deliver impressive performance in isolation, they have also fragmented the alignment landscape, increasing workflow complexity and reducing accessibility for non-specialists. Because different seeding strategies require dedicated algorithms and internal data structures to achieve optimal performance and efficiency, it remains unclear whether the diverse optimizations demanded by disparate alignment tasks can be integrated within a single algorithmic framework (see the Background section). Consequently, developing a truly multi-purpose aligner that maintains extensibility while delivering consistently high performance and computational efficiency remains an open challenge in both theory and practice.

Here we present the Versatile Alignment Toolkit (VAT), a unified algorithmic framework for biological sequence alignment. VAT is designed to address three theoretical and practical goals: (1) to bridge the gap between a unified theoretical framework for sequence alignment and the proliferation of task-specific aligners in practice, demonstrating that a single high-performance, multi-purpose, and computationally efficient aligner is feasible; (2) to simplify the analysis of emerging biological sequence data, which may differ in alphabet, length, identity, structure, and scale as technologies continue to evolve; and (3) to streamline the construction of bioinformatic pipelines that integrate multiple alignment tasks. To achieve these goals, VAT enables run-time parameterization of alignment settings without reindexing the reference database. It introduces an asymmetric multi-view indexing scheme that encodes multiple seeding strategies within a single reference index, allowing users to switch seeding schemes dynamically. VAT further leverages longest common prefix (LCP) information to retrieve seed locations for variable seed lengths, eliminating the need to rebuild indexes when seed-length parameters change. A parameterized seed-chaining algorithm optimizes seed arrangements for specific alignment objectives, an essential capability for tasks such as split-read mapping and whole-genome alignment. In addition, VAT incorporates in-register bitonic sorting and on-the-fly streaming-based seed matching to reduce computational overhead and memory usage during index construction and seeding. Collectively, these design choices enable VAT to flexibly accommodate sequence data ranging from short and long reads to entire genomes through simple parameter tuning rather than algorithmic modification.

We benchmark VAT with state-of-the-art methods on short- and long-read mapping, DNA and protein homology search, and whole-genome alignments, representing sequence data variability across all dimensions. As a single multi-purpose aligner, VAT demonstrated superior or on-par performances compared to the specialized aligners in their respective fields. By decoupling data representation from alignment logic, VAT achieves both high performance and broad applicability, establishing a generalizable foundation for future sequence analysis tools. VAT is freely available for academic use at https://github.com/xuan13hao/VAT.

## 2. Background

In this section, we provide a technical overview of major seeding and indexing strategies, which are central determinants of an aligner’s sensitivity, specificity, and computational performance. The core intuition behind the seed-and-extend framework is to restrict detailed alignment to sequence pairs that share regions of high local similarity^21-23,45^. Classical approaches achieve this by decomposing both reference and query sequences into fixed-length substrings, or *k*-mers, and initiating alignment only when at least one *k*-mer is shared between the two sequences^46,47^. Varying *k m*odulates performance: shorter *k* -mers increase sensitivity but risk spurious matches, whereas longer *k*-mers improve specificity and speed at the cost of reduced sensitivity. More refined strategies connect the number of shared *k* -mers to the estimated sequence similarity, requiring multiple *k* -mer matches to trigger extension and thereby reducing opportunistic hits between unrelated sequences^28^. However, when aligning highly similar sequences, the use of densely overlapping *k* -mers produces large numbers of redundant matches, thereby reducing computational efficiency. To mitigate this, minimizers were introduced to reduce redundant *k* -mers: a minimizer is the lexicographically smallest *k*-mer within an *h*-mer window (*h* > *k*), such that only a single representative *k*-mer is selected for each window^30,34,48^. This reduces the total number of seed candidates while preserving the ability to detect high-identity alignments. Overall, both *k*-mer–based and minimizer-based strategies are conceptually simple, efficient, and highly effective for mapping sequences with substantial similarity (e.g., high-throughput sequencing mapping).

However, the strategies described above may perform poorly when aligning low-similarity sequences, which may be too divergent to share long exact matches. One straightforward solution is to relax the requirement for exact matching by allowing inexact *k* -mer matches whose similarity scores exceed a threshold (e.g., based on BLOSUM or PAM substitution matrices)^49,50^. Although highly sensitive in principle, this approach is computationally inefficient because evaluating inexact matches is far more expensive than detecting exact ones. Two alternative strategies circumvent this limitation by implicitly tolerating mismatches while still computing only exact matches. The first is the spaced-seed (seed-pattern) approach, which applies a mask to each *k*-mer, requiring exact matches only at unmasked positions^29,51^. For example, under the pattern “101” (where “1” indicates retained positions and “0” indicates masked positions), the 3-mers “ABC” and “ADC” both map to “AC,” thereby forming a match despite a mismatch at the middle position. The second strategy uses a reduced alphabet, in which residues with similar physicochemical properties are grouped into the same symbols^35,52,53^. For instance, lysine (K) and arginine (R) collapse into a single letter, allowing “KFG” and “RFG” to match exactly in the reduced alphabet even though they differ in the original one. These inexact-tolerant seeding schemes are commonly employed in homology search and other applications involving sequences with low evolutionary identity.

Most aligners precompute *k*-mer information for the reference database during an indexing step to reduce the computational cost of seed matching^10,31,54^. Classical approaches rely on hash tables, in which *k*-mers serve as keys and their genomic positions as values; query-derived *k*-mers are then used to probe this hash structure during seed matching^32,34^. Other aligners store the same information in a lexicographically sorted *k*-mer table, using binary search–like procedures to locate matching entries^55,56^. Although simple and effective, both approaches share two key limitations: (1) the index cannot be reused when the seed length *k* changes, restricting runtime parameter flexibility; and (2) querying the table can be slow, especially for large reference databases. To address these limitations, more advanced full-text indexing structures such as suffix trees^57,58^, suffix arrays^38,40,59,60^, and the Burrows–Wheeler Transform (BWT) were introduced^38,42,61^. These data structures compactly represent all suffixes of the indexed sequence and support highly efficient exact matching for arbitrary seed lengths, enabling true runtime seed-length parameterization. However, they often require substantial memory and loading them in full can be challenging on lower-end hardware. To mitigate these limitations, a class of run-length compressed BWT-based full-text indexes has been developed, including the r-index and move-structure-based approaches. These methods exploit the repetitiveness of genomic and pangenomic data by scaling with the number of runs in the BWT rather than the total sequence length, substantially reducing memory usage while retaining the flexibility of full-text search^62,63^. Orthogonally to index compression, the double-indexing strategy avoids holding large reference indexes in memory altogether^44^. Instead, both the reference and the query are converted into lexicographically sorted *k*-mer tables: the reference table is precomputed, whereas the query table is generated at runtime. The two tables are then streamed simultaneously from disk, and seed matches are detected using a merge-join algorithm. Although this strategy introduces the overhead of indexing the query, it gains efficiency by eliminating the need to load large reference indexes into RAM and by exploiting the inherent speed of merge-join operations. When implemented carefully, double indexing provides considerable performance benefits over approaches that index only the reference database.

## 3. Methods

### 3.1. Overall design

The design principle underlying VAT is to integrate all major seeding schemes reviewed above and to permit run-time adjustment of seed length without reindexing the reference database. This flexibility allows VAT to be easily parameterized and adapted to diverse alignment tasks across scales and applications. Specifically, VAT follows a seed-and-extend architecture that generalizes the canonical sequence alignment paradigm (Figure 1a). It first constructs asymmetric multi-view indexes for the reference and query datasets (indexing), then identifies seed matches between them (seeding). These seeds are subsequently extended both ungapped and gapped (extension), and the resulting fragments are chained through dynamic programming to determine an optimal alignment layout (chaining). Remaining gaps between chained regions are resolved using a modified Needleman–Wunsch algorithm to yield the final alignment (gap-filling). Within this architecture, VAT introduces five key innovations. Some are designed to expand generalizability across diverse alignment tasks (Figure 1a, light blue boxes), and others focused on achieving high computational efficiency (Figure 1a, green boxes). The complete VAT algorithm is detailed in the Supplementary Methods.

**Figure 1:**
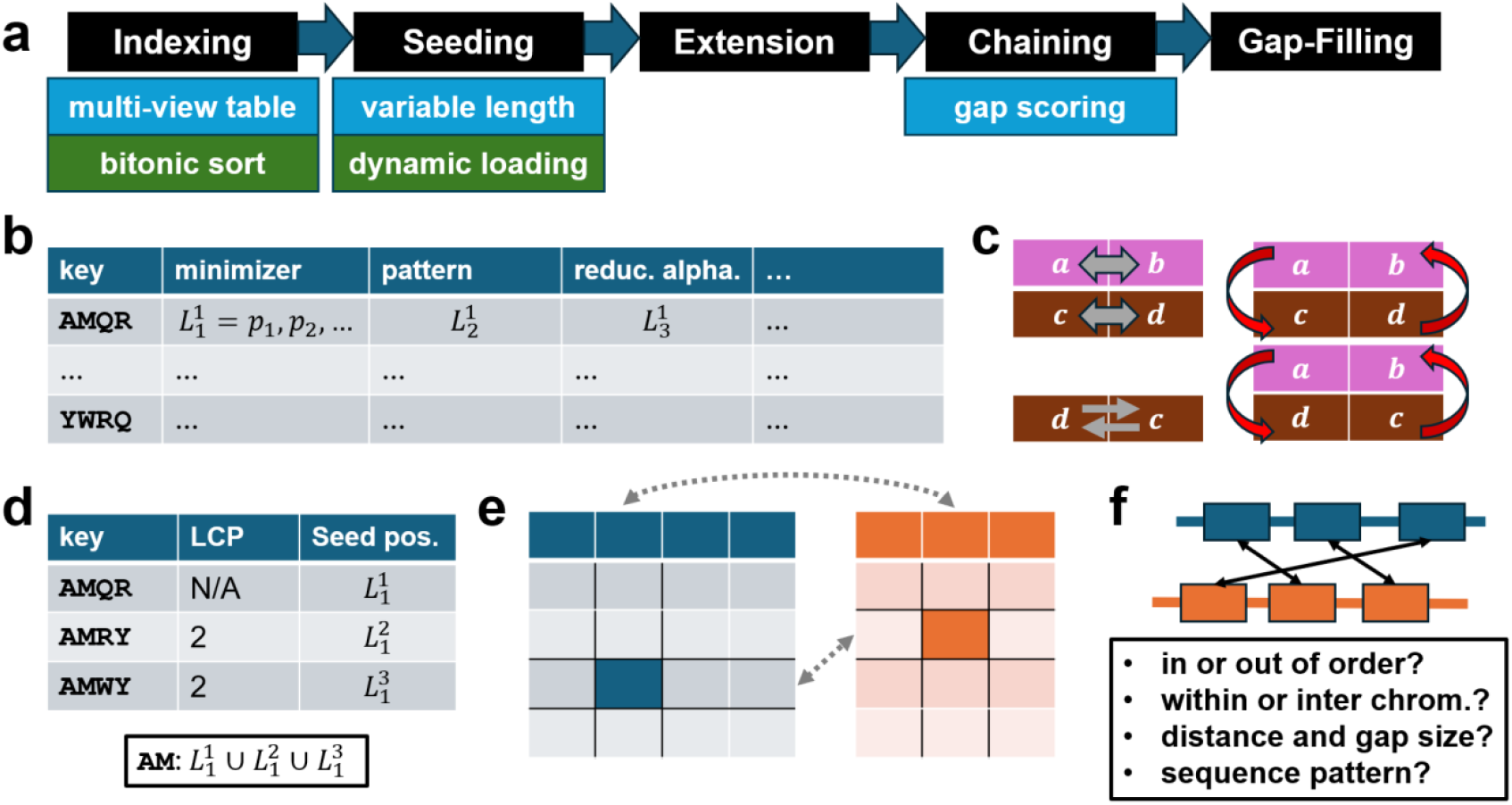
Overview of VAT’s unified algorithmic framework and key design components. (a) Schematic of the major components of VAT. Light blue boxes highlight VAT-specific design elements that enable broad generalizability across alignment tasks, whereas green boxes indicate optimizations that enhance computational efficiency. (b) Illustration of VAT’s multi-view index table. Each row corresponds to a unique key (short string), and each column represents a distinct seeding scheme; individual table entries store the genomic positions at which the key appears under the corresponding seeding pattern. (c) Toy example of in-register bitonic sorting. Each pair of pink or brown blocks represents the two data elements stored within a single register. Gray arrows denote in-register operations, and red arrows denote cross-register comparisons used to complete the bitonic sorting sequence. (d) Illustration of how VAT uses longest common prefix (LCP) values to compute genomic positions for shorter seeds without reindexing. In this example, the genomic positions for the 2-mer “AM” are obtained by taking the union of entries within a consecutive table range whose minimum LCP value is 2. (e) Overview of VAT’s dynamic seeding strategy. The indigo and orange tables represent reference and query indexes, respectively. Only table entries matched in both row and column dimensions are loaded into memory, thereby minimizing memory usage. (f) Summary of the task-specific information incorporated during seed chaining, including the ordering of the seeds, their positional relationships, and anticipated sequence patterns.

### 3.2. Asymmetric multi-view indexing

VAT constructs asymmetric multi-view indexes for the reference dataset *R* and the query dataset *Q*. In a multi-view table (Figure 1b), each row is keyed by a unique short sequence *s*_*i*_, and each column corresponds to a distinct seeding scheme *T*_*j*_. The table is sorted lexicographically by its keys (*s*_*i*_). Each entry in the table stores a list of genomic positions 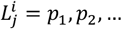,representing all locations at which the key sequence *s*_*i*_ occurs under seeding scheme *T*_*j*_. In other words, if *s*_*i*_ = *T*_*j*_(Genome(*p*_*x*_)) for some positions *p*_*x*_, *p*_*x*_ is appended to the list 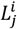. This compact representation encodes the reference under multiple seeding strategies simultaneously, enabling VAT to switch seamlessly among them. VAT currently supports exact-match^37,64-66^, minimizer^30,48^, spaced-pattern^29,51^, and reduced-alphabet seeding schemes^35,52,53^, and additional schemes can be incorporated without modifying the overall framework. During index construction, all supported seeding schemes are applied to the reference, whereas the query index includes only those schemes specified by the user at runtime. This asymmetric multi-view indexing design enables VAT to flexibly adapt to heterogeneous datasets and alignment objectives without rebuilding the index, thereby providing both generality and computational efficiency. Moreover, it allows the construction of a single, reusable reference index that supports multiple alignment tasks. For example, a single human genome index can be used seamlessly for short- and long-read mapping, homology search, and whole-genome alignment against the same reference, substantially simplifying complex bioinformatic workflows.

### 3.3. In-register bitonic string sorting

To efficiently sort the keys in the multi-view tables, VAT employs in-register bitonic sorting, which exploits Single-Instruction Multiple-Data (SIMD) operations for parallel string comparison^67-69^ (Figure 1c). Specifically, VAT processes four strings (*a, b, c*, and *d*) simultaneously using two 256-bit registers (*X*, Figure 1c pink blocks; and *Y*, Figure 1c brown blocks), each holding a pair of 128-bit strings. The algorithm first compares string pairs within each register (*a* with *b* and *c* with *d*), then performs cross-register comparisons in parallel to compare *a* with *c* and *b* with *d*. Next, it swaps *c* and *d* in register *Y*, and performs cross-register comparison again to compare *a* with *d* and *b* with *c*. In this way, it evaluates all pairwise orderings among the four strings. In this way, VAT sorts four data elements with only three in-register steps (two comparisons and one swap) and two cross-register comparisons. This hardware-efficient implementation minimizes memory access and instruction cycles, enabling rapid lexicographic sorting of large sequence indexes and substantially improving indexing speed.

### 3.4. Variable-length seeding information retrieval

VAT leverages the longest common prefix (LCP) information to support seed-length parameterization without reindexing (Figure 1d). VAT computes the LCP value for each key, defined as the length of the shared prefix between the key and its preceding key in lexicographic order^61^, and stores these values in a dedicated column of the multi-view index table. Within any contiguous range of entries, the effective common prefix length (e.g., the shared common prefix “AM” in Figure 1d) is determined by the minimum LCP value across the range. The genomic positions of the shared common prefix can be obtained by taking the union of the corresponding positional lists (Figure 1d,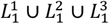). By dynamically adjusting the LCP threshold, VAT aligns seeds consistent with user-specified lengths, achieving flexible and memory-efficient seed matching across diverse datasets.

### 3.5. Memory-efficient and dynamic seed matching

VAT streams both the reference and query indexes simultaneously from disk (Figure 1e), matching seeds using the merge-join algorithm without loading the entire indexes into memory. Only when both the key strings (Figure 1e, matched rows) and the seeding schemes (Figure 1e, matched columns) match does VAT loads the corresponding genomic-position lists (Figure 1e, entries with solid colors) into main memory. This selective loading strategy ensures that memory is used only for entries relevant to the current seed match. As a result, VAT maintains a modest memory footprint despite operating on a more comprehensive multi-view index structure.

### 3.6. Flexible seed chaining via parameterized gap cost

VAT adopts a dynamic programming–based chaining algorithm similar to those used in Minimap2^33^ and CLAN^70^. For each pair of candidate seeds, VAT computes a chaining gap penalty defined as a weighted function of several factors: the relative ordering of the seeds, whether they occur on the same chromosome, the genomic distance between them, the size of the intervening gap, and the presence of expected sequence patterns (Figure 1f). The weights for these factors are task-specific. For example, in RNA-seq read mapping, valid seed chains must lie on the same chromosome, be separated by distances consistent with typical intron lengths, and exhibit canonical donor–acceptor motifs at splice junctions^71-74^. In contrast, for CLASH data, seeds originating from interacting miRNAs and mRNAs may reside on different chromosomes, and VAT’s chaining logic accommodates such cross-chromosomal pairings. This task-aware scoring formulation enables VAT to generalize across diverse biological contexts while maintaining biologically meaningful alignment structure.

## 4. Results

### 4.1. Benchmark experiment design

We benchmarked VAT against state-of-the-art aligners across four major application domains: short-read mapping, long-read mapping, homology search, and whole-genome alignment. All task-specific VAT parameters, detailed descriptions of the benchmark datasets, and the definitions of all reported performance metrics are provided in the Supplementary Methods.

### 4.2. High-throughput sequencing read mapping

We first evaluated VAT in high-throughput sequencing (HTS) read mapping^75-77^, encompassing both short reads from next-generation sequencing (NGS) and long reads from third-generation sequencing (TGS) platforms. HTS read mapping tests an aligner’s ability to efficiently process vast volumes of highly similar sequences^78,79^—a critical capability for modern genomic analysis.

#### 4.2.1. Next-generation sequencing read mapping

Next-generation sequencing (NGS) reads, such as those generated by Illumina platforms, are typically short (≈100-150 bp), highly accurate (<0.1% error rate), and produced in massive volumes^76,80,81^. Depending on library preparation, reads may derive from contiguous genomic regions (e.g., whole-genome or exome sequencing) or from discontinuous regions (e.g., RNA-seq or Hi-C)^82,83^. To assess VAT’s performance in contiguous read mapping, we analyzed one simulated and four real datasets spanning diverse applications: human whole-genome sequencing (WGS)^84,85^, chromatin immunoprecipitation sequencing (ChIP-seq)^86^, assay for transposase-accessible chromatin using sequencing (ATAC-seq)^87^, and 16S rRNA sequencing of the human gut microbiome (16S microbiome)^88^. We further evaluated VAT’s generalizability across five representative species—*Caenorhabditis elegans, Drosophila melanogaster, Mus musculus, Oryza sativa, and Arabidopsis thaliana* (Supplementary Table S1). We compared VAT with leading short-read aligners representing diverse algorithmic architectures, including BBMAP^89^, BLASR^90^, Bowtie2^91^, BWA-MEM^41^, HISAT2^71,72^, Minimap2^33,34^, SOAP2^92^, STAR^39^, segemehl^93^, Tophat2^94,95^ and Subread^64^. All programs were executed using recommended parameters (Supplementary Table S2).

Figure 2a–e and Supplementary Tables S3–S7 summarize the contiguous read mapping performance of VAT and other benchmarked aligners across simulated and experimental datasets. On simulated reads (Figure 2a), VAT aligned 99.8% of reads with 98.9% accuracy, matching the highest mapping rate achieved by BLASR (100%) while exceeding its accuracy (97.1%). HISAT2 attained slightly higher accuracy (99.1%) but with a lower mapping rate (97%). Across real datasets (Figure 2b–e), VAT achieved the highest mapping rate for WGS, ranked second for ChIP-seq (99.0% vs. BWA MEM’s 99.37%) and ATAC-seq (99.7% vs. BLASR’s 100%), and third for 16S microbiome data (95.3% vs. BLASR’s 96.9% and BWA MEM’s 95.5%). VAT maintained consistently high mapping rates across multiple non-human species, ranking second overall (99.8% vs. STAR’s 100%; Supplementary Tables S8–S12). Despite its high alignment sensitivity, VAT remained computationally efficient (Figure 2f–j; Supplementary Tables S3–S12). Averaged across all datasets, VAT and BLASR achieved comparable mapping rates (99.1%), yet VAT operated ≈49-fold faster. Although HISAT2 was marginally faster than VAT (2.5%), its average mapping rate was substantially lower (84.3%). Collectively, these results highlight VAT’s unique balance of alignment sensitivity, accuracy, and computational efficiency in mapping contiguous NGS reads across diverse genomic contexts.

**Figure 2:**
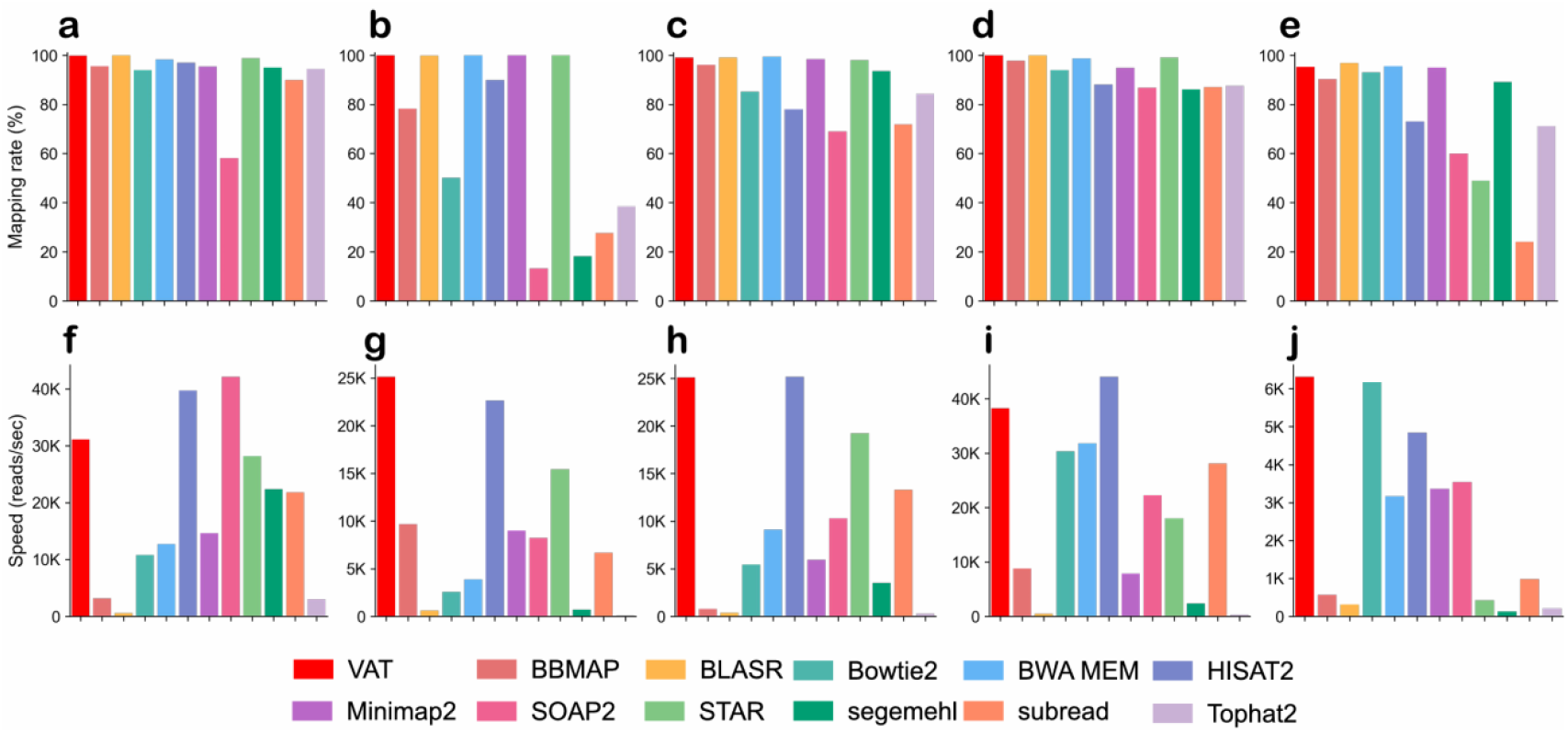
Contiguous short-read mapping performance of VAT and benchmarked aligners. Panels (a–e) show the percentage of reads successfully mapped for the simulated WGS, real WGS, ChIP-seq, ATAC-seq, and 16S rRNA gut microbiome datasets, respectively. Panels (f–j) display the corresponding mapping speeds, measured as reads aligned per second.

For split-read mapping, we evaluated VAT using one simulated RNA-seq dataset and four experimental datasets derived from RNA-seq, circular RNA sequencing (circRNA-seq)^96-98^, cross-linking ligation and sequencing of hybrids (CLASH)^70,99^, and high-throughput chromatin conformation capture (Hi-C)^100,101^. These datasets represent distinct biological contexts characterized by specific split-read patterns: splice junctions in RNA-seq, back-splicing in circRNA-seq, chimeric microRNA–mRNA interactions in CLASH, and topologically associating domains (TADs) in Hi-C. We further included RNA-seq data from five non-human model organisms to assess cross-species generalizability. Details regarding all benchmark datasets are available in Supplementary Table S13. Ground-truth annotations were defined according to established resources: Ensembl GRCh38^102^ for known splice junctions, circAtlas for circular RNAs^103^, miRBase^104^ and TargetScan^105^ for CLASH interactions, and experimentally defined TADs^106^ for Hi-C (Supplementary Methods). VAT was benchmarked against BBMAP, BLASR, Bowtie2, BWA-MEM, HISAT2, Minimap2, SOAP2, STAR, segemehl, TopHat2, and Subread, each executed with recommended parameters (Supplementary Table S14).

We evaluated three primary metrics across all tools—overall mapping rate, split-read mapping rate, and recovery of known split events. On the simulated RNA-seq dataset (Figure 3a,f; Supplementary Tables S15) and the real circRNA-seq dataset (Figure 3c,h; Supplementary Tables S16), VAT ranked first in all three metrics. On the real RNA-seq dataset (Figure 3b,g), VAT achieved the highest overall mapping rate and known-event recovery, and ranked second in split-read mapping rate (36.5% vs. STAR’s 37.2%; Supplementary Table S17). For the CLASH dataset (Figure 3d,i; Supplementary Tables S18), VAT ranked second in overall mapping rate (97.2% vs. Bowtie2’s 98.2%) and first in both split-read mapping rate and recovery of known miRNA–mRNA chimeric junctions. On the Hi-C dataset (Figure 3e,j; Supplementary Tables S19), VAT ranked third in split-read mapping rate (4.1%, compared with segemehl’s 6.4% and STAR’s 6.1%) but achieved the highest recovery of biologically validated split events (VAT 32.3%, segemehl 26.9%, STAR 18.2%), indicating superior alignment precision. Across RNA-seq datasets from non-human model organisms (Supplementary Tables S20–S24), VAT ranked first for all metrics except for *O. sativa*, where it placed second in overall mapping rate (99.2% vs. STAR’s 99.4%). Importantly, VAT consistently delivered these gains while being the fastest aligner across all split-read datasets (Figure 3k–o).

**Figure 3:**
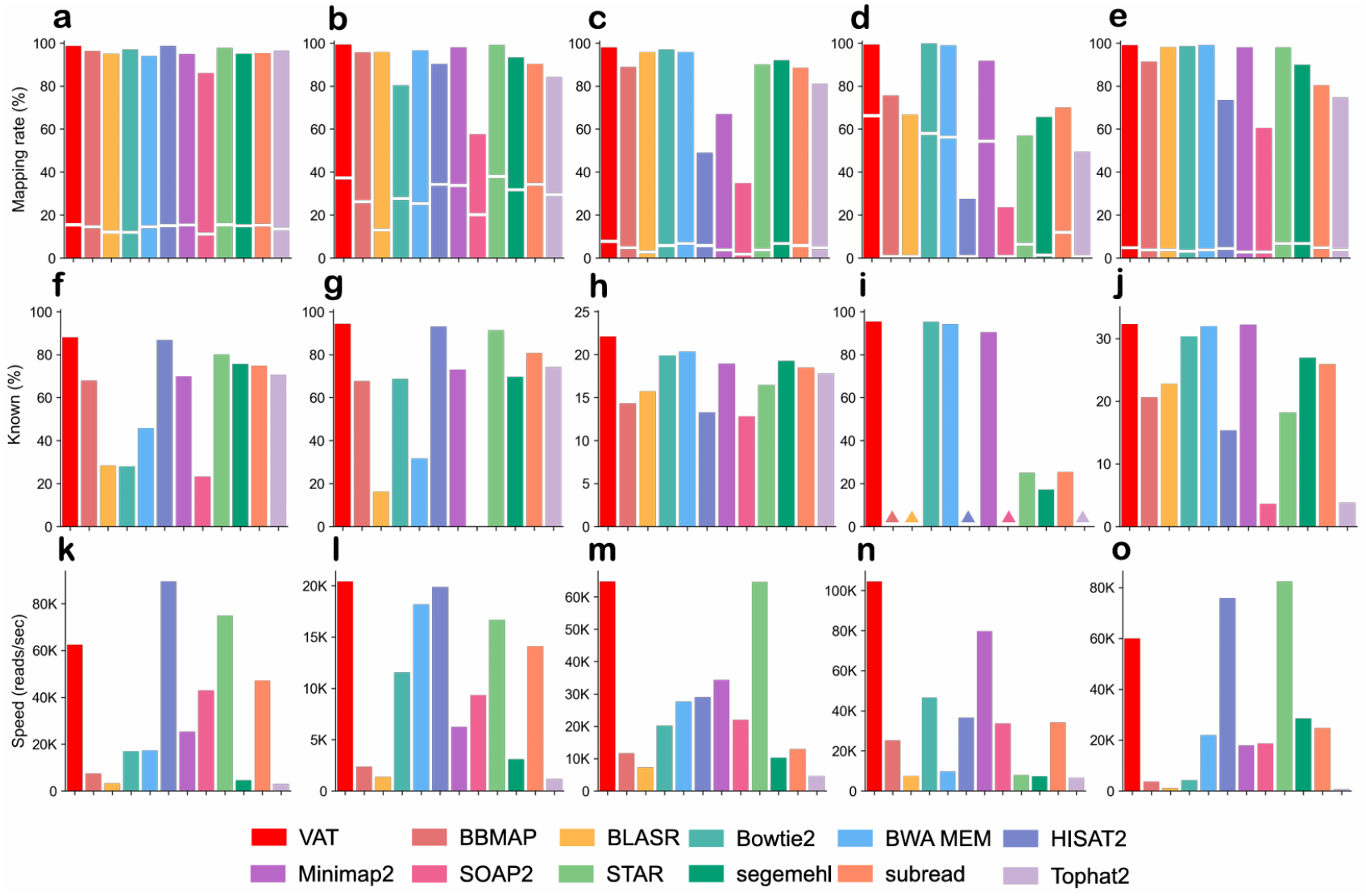
Contiguous short-read mapping performance of VAT and benchmarked aligners. Panels (a–e) show the percentage of reads successfully mapped for the simulated RNA-seq, RNA-seq, circRNA-seq, CLASH, and Hi-C datasets, respectively. Bars above the breaks represent the proportion of mapped contiguous reads,while those below indicate mapped split reads. Panels (f–j) show the proportion of known split events. Panels (k–o) display the corresponding mapping speeds, measured as reads aligned per second.

#### 4.2.2. Third-generation sequencing read mapping

Third-generation sequencing (TGS) reads are typically long (often >1 kb), moderately accurate with platform-dependent error rates, and produced at lower throughput than NGS^7,107,108^. Like NGS reads, they may originate from contiguous genomic regions (e.g., WGS) or from split regions (e.g., RNA-seq). Mapping TGS reads demands algorithms capable of handling long sequences with reduced identity, particularly when multiple splice junctions present within a single read. To assess performance under stringent conditions, we performed all benchmarks without prior error correction, thereby evaluating each aligner in a worst-case scenario.

For contiguous long-read mapping, we evaluated VAT on one simulated WGS dataset and four experimental datasets: human WGS, ChIP-seq, ATAC-seq, and 16S rRNA human gut microbiome (Supplementary Table S25). Additional long-read datasets from five representative model organisms were also analyzed (Supplementary Table S25). VAT was benchmarked against six state-of-the-art long-read aligners—BLASR^90^, GMAP^109^, GraphMap2^110^, ngmlr^111^, Minimap2^33^, and STARlong^39,83^, each executed with recommended parameters (Supplementary Table S26).

Figure 4a–e and Supplementary Tables S27–S31 summarize the performance of VAT and benchmarked long-read aligners on contiguous TGS datasets. On the simulated WGS dataset (Figure 4a; Supplementary Table S27), VAT achieved the highest mapping rate (96.8%) and accuracy (98.6%). This trend was consistent across all experimental datasets, including those derived from human and multiple model organisms (Figure 4b-e; Supplementary Tables S28–S36). Notably, VAT also exhibited the highest computational efficiency (Figure 4f–j), operating approximately 60% faster than the next fastest aligner, Minimap2, while maintaining superior alignment accuracy. These results highlight VAT’s ability to balance speed, sensitivity, and precision in long-read mapping across diverse sequencing contexts.

**Figure 4:**
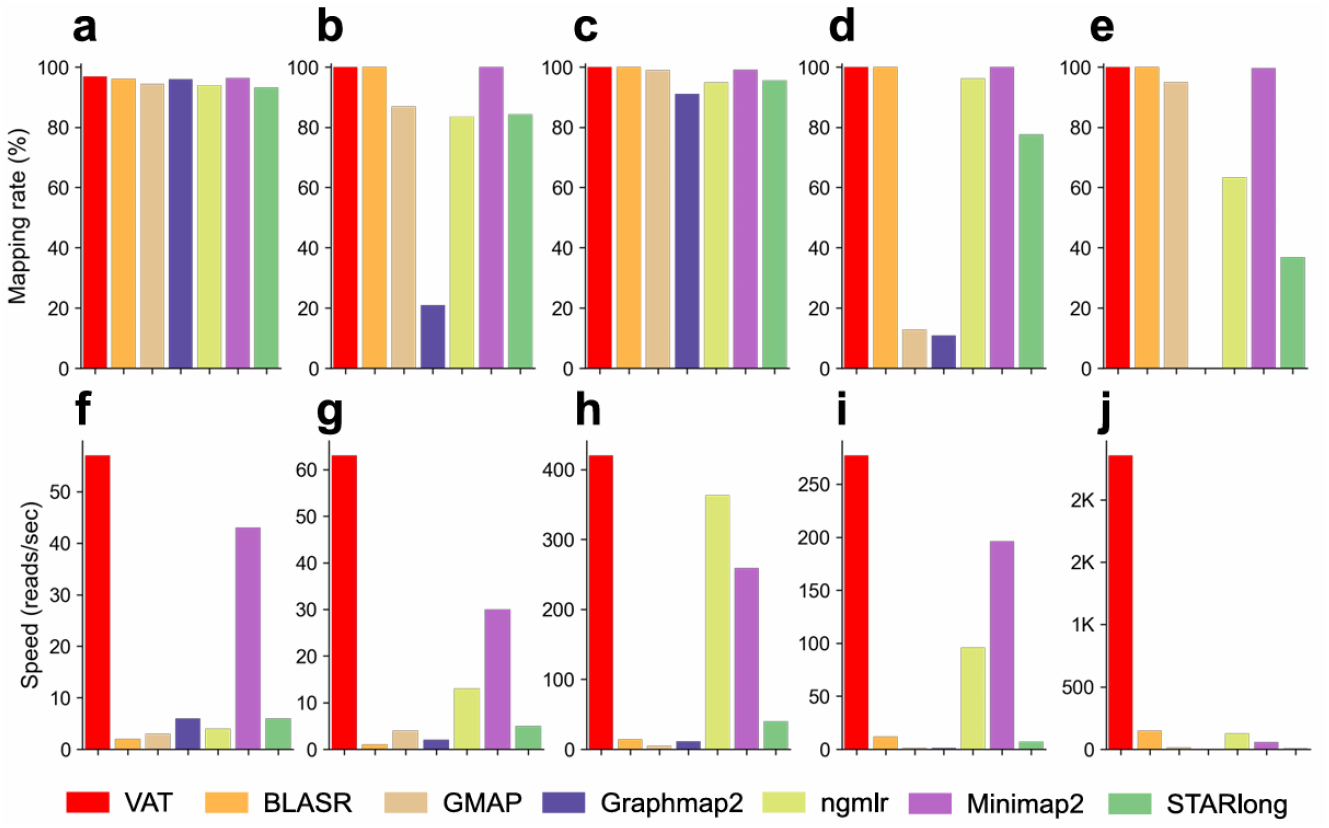
Contiguous long-read mapping performance of VAT and benchmarked aligners. Panels (a–e) show the percentage of reads successfully mapped for the simulated WGS, WGS, ChIP-seq, ATAC-seq, and 16S rRNA gut microbiome datasets, respectively. Panels (f–j) display the corresponding mapping speeds, measured as reads aligned per second.

For split-read mapping of TGS data, we evaluated VAT on one simulated and one real human RNA-seq dataset, together with five RNA-seq datasets from representative model organisms (Supplementary Table S37). These datasets assess aligner performance on long reads spanning multiple splice junctions, a key challenge in transcript-level analysis. VAT was benchmarked against six state-of-the-art long-read aligners—BLASR, GMAP, GraphMap2, ngmlr, Minimap2, and STARlong—each executed with recommended parameters (Supplementary Table S38).

Figure 5a–b and Supplementary Tables S39–S40 summarize the split long-read mapping performance of VAT and benchmarked aligners. Across both datasets, VAT achieved the highest overall mapping rates (98.0% and 96.8%) and the highest split-read mapping rates (75.1% and 70.8%). Figure 5c–d show that VAT produced the largest proportion of split reads correctly spanning known splice sites (83.4% and 84.5%). VAT also demonstrated superior computational efficiency (Figure 5e–f), ranking as the fastest aligner among all tested tools. The same trend was observed across datasets from five representative model organisms (Supplementary Tables S41–S45), underscoring VAT’s robust and scalable performance in split-read long-read mapping.

**Figure 5:**
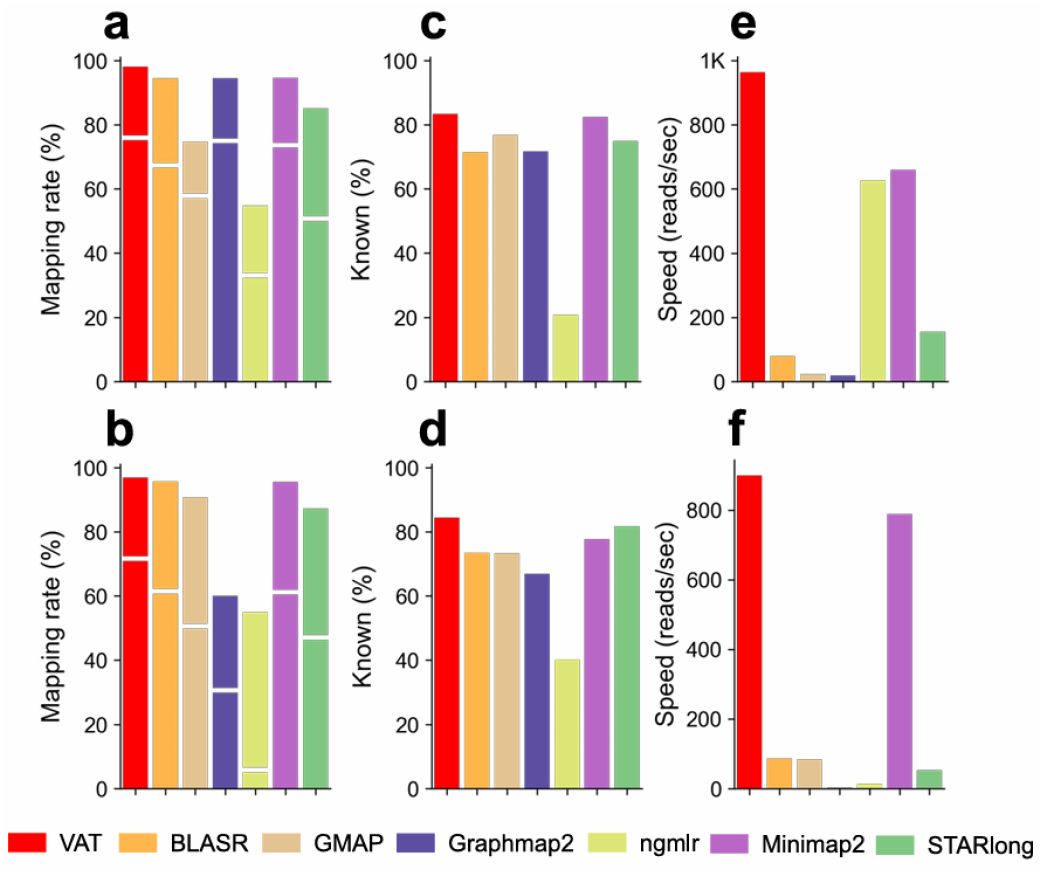
Split long-read mapping performance of VAT and benchmarked aligners. Panels (a–b) show the mapping rate for the simulated and real RNA-seq datasets, respectively. In each panel, bars above the axis break represent the proportion of contiguously mapped reads, whereas bars below represent split-read alignments. Panels (c–d) display the fraction of split reads correctly spanning known splice sites for the simulated and real datasets. Panels (e–f) present the corresponding mapping speeds, measured as reads aligned per second.

### 4.3. Homology search

We next evaluated VAT in the context of homologous sequence search across protein and DNA datasets, a task that challenges an aligner’s ability to detect low-identity relationships (often as low as 20%)^45,112,113^. For protein homology, we constructed an evaluation dataset from Pfam^114^, using its curated family classifications as ground-truth labels. DNA-level homology was assessed using Homologous Gene Database (HGD)^115^. Within each dataset, we performed all-against-all pairwise alignments. VAT was benchmarked against BLAST^21^, MMseqs2^116^, DIAMOND^43^, and RAPSearch2^117^ for protein homology.

Because VAT, DIAMOND, and MMseqs2 each provide distinct fast and sensitive variants, these modes were benchmarked individually. VAT was further benchmarked with BLAST and pblat^118,119^ for DNA homology. All programs were invoked using recommended parameters (Supplementary Tables S46-S47).

Figure 6a and Supplementary Table S48 summarize the performance of VAT and benchmarked tools on protein homology search. Among aligners configured for maximum sensitivity (BLASTP, DIAMOND– ultra-sensitive, MMseqs2–s7.5, and VAT–sensitive), BLASTP achieved the highest F1 score (78.2%), followed closely by VAT–sensitive (77.3%), DIAMOND–ultra-sensitive (76.8%), and MMseqs2–s7.5 (76.2%). At this performance tier, VAT–sensitive operated ∼10% faster than DIAMOND–ultra-sensitive, ∼50% faster than MMseqs2–s7.5, and 7.5-fold faster than BLASTP (Figure 6b). Among aligners optimized for speed (VAT–fast, DIAMOND–fast, and MMseqs2–s1), VAT–fast delivered the highest F1 score (52.8%), substantially outperforming DIAMOND–fast (31.4%) and MMseqs2–s1 (35.8%) (Figure 6a). VAT–fast and DIAMOND–fast exhibited comparable runtime performance, both approximately 30% faster than MMseqs2–s1 (Figure 6b). RAPSearch2 showed intermediate performance, with an F1 score of 62.6% and speed falling between the sensitive and fast tiers (Figure 6a,b).

**Figure 6:**
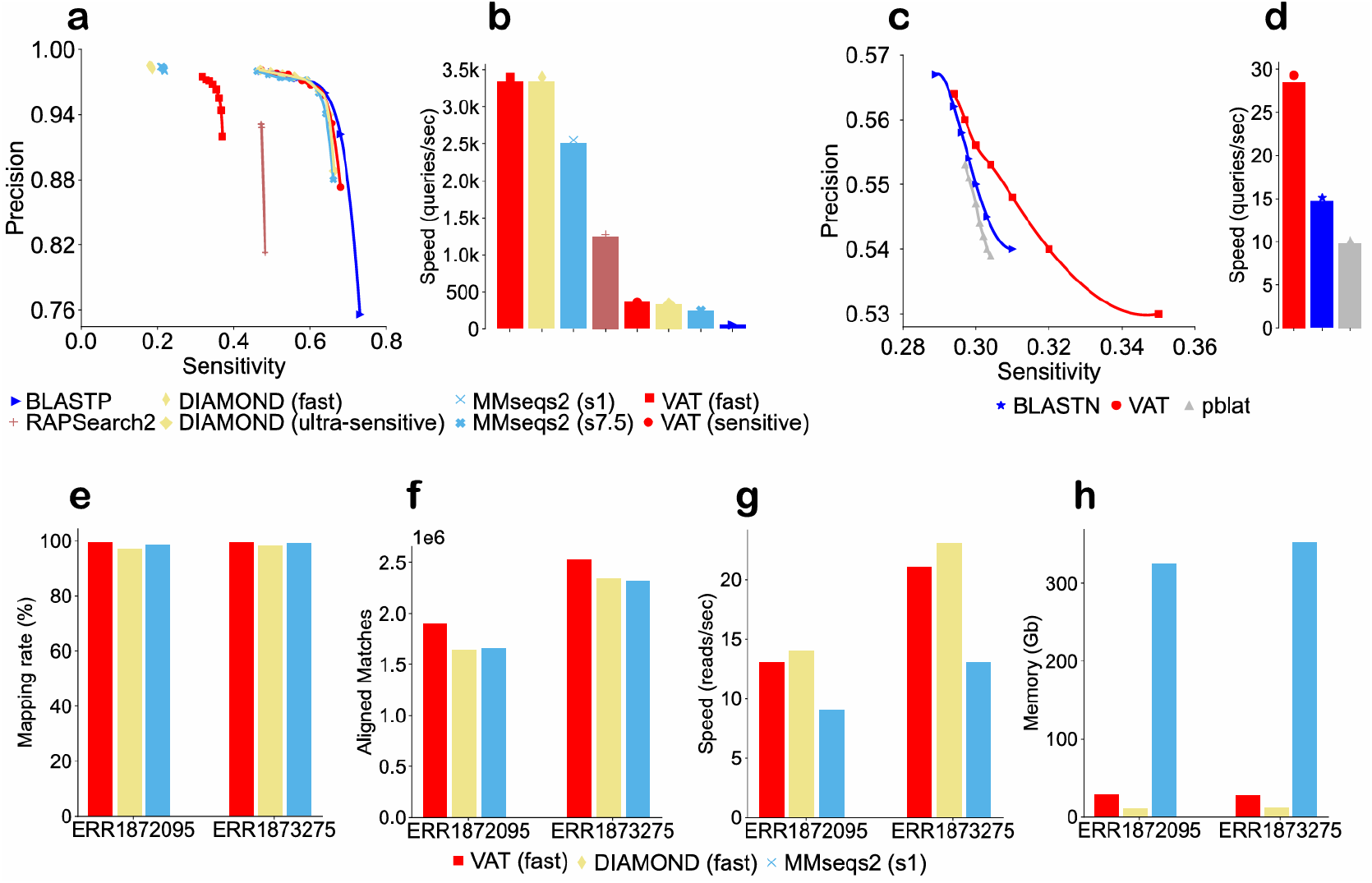
Homology search performance of VAT and benchmarked aligners. (a) Receiver operating characteristic (ROC) curves for VAT and benchmarked tools on protein homology search. (b) Corresponding running times for protein homology search. (c) ROC curves for VAT and benchmarked tools on DNA homology search. (d) Corresponding running times for DNA homology search. (e) Alignment rates for VAT-fast, DIAMOND-fast, and MMseqs2-s1 when aligning two soil metagenomic datasets against the NR database. (f) Number of homologs detected from the NR database. (g) and (h) Corresponding alignment speeds and memory consumption.

Figure 6c and Supplementary Table S49 summarize the performance of VAT and benchmarked tools on DNA homology search. VAT achieved the highest overall F1 score (42.2%), outperforming BLASTN (39.4%) and pblat (38.9%). Relative to BLASTN, VAT attained higher sensitivity (35% vs. 31%) with only a modest reduction in precision (53% vs. 54%) while operating at twice the speed (Figure 6d). Consistent with observations from the protein homology search benchmarks, many of the false positives identified by VAT exhibited substantial sequence similarity when examined across multiple alignment methods, suggesting that VAT captures biologically plausible relationships that may fall near the boundary of existing reference-based classifications.

In addition to standard protein–protein and DNA–DNA homology search, VAT also supports cross-domain alignments between proteins and translated nucleotide sequences, analogous to BLASTX and TBLASTN. To assess performance in this setting, we benchmarked VAT-fast, DIAMOND-fast, and MMseqs2-s1 by searching two soil metagenomic datasets (Supplementary Table S50) against the full NR database. VAT-fast achieved the highest alignment rate (99.3%), followed by MMseqs2-s1 (98.7%) and DIAMOND-fast (97.5%) (Figure 6e; Supplementary Table S50). VAT-fast also detected 11.8% more homologous sequences than DIAMOND-fast and 12.0% more than MMseqs2-s1 (Figure 6f; Supplementary Table S50). In terms of computational efficiency, VAT-fast was 7.9% slower than DIAMOND-fast but 53.0% faster than MMseqs2-s1 (Figure 6g; Supplementary Table S50). VAT-fast consumed 63.2% more memory than DIAMOND-fast, yet both tools remained substantially more memory-efficient, by nearly an order of magnitude, than MMseqs2-s1 (Figure 6h; Supplementary Table S50).

### 4.4. Whole-genome alignment

Whole-genome alignment (WGA) presents a stringent challenge for alignment tools due to the presence of extremely long sequences and extensive structural variation, including segmental duplications, deletions, and inversions. To assess VAT under these conditions, we compared it with MUMmer4^57^ by aligning the human and chimpanzee genomes as well as between two *Arabidopsis* genomes (Supplementary Table S51). Parameter settings and full command lines for both tools are provided in Supplementary Table S52.

Figure 7a and Supplementary Table S53 summarize the human-chimpanzee WGA coverage produced by VAT and MUMmer4 across a range of identity thresholds. At each threshold, VAT achieved 0.3–2.2% higher coverage than MUMmer4, reaching 87.8% coverage at 90% identity. A similar trend was observed for alignments between the two *Arabidopsis* genomes (Supplementary Table S54). Figure 7b shows the WGA dot plot for the human and chimpanzee X chromosome, a region known to be highly conserved. VAT recovered all alignments detected by MUMmer4 and additionally identified several inversion events not reported by MUMmer4. For the Y chromosome, which is substantially more divergent between human and chimpanzee, VAT again captured all MUMmer4 alignments while revealing a markedly larger number of homologous regions (Figure 7c). All reported alignments in Figure 7b,c are longer than 6K bp and have sequence identity higher than 90.0%. VAT completed the human–chimpanzee whole-genome alignment in 43 minutes, making it more than three times faster than MUMmer4 (Supplementary Table S55).

**Figure 7:**
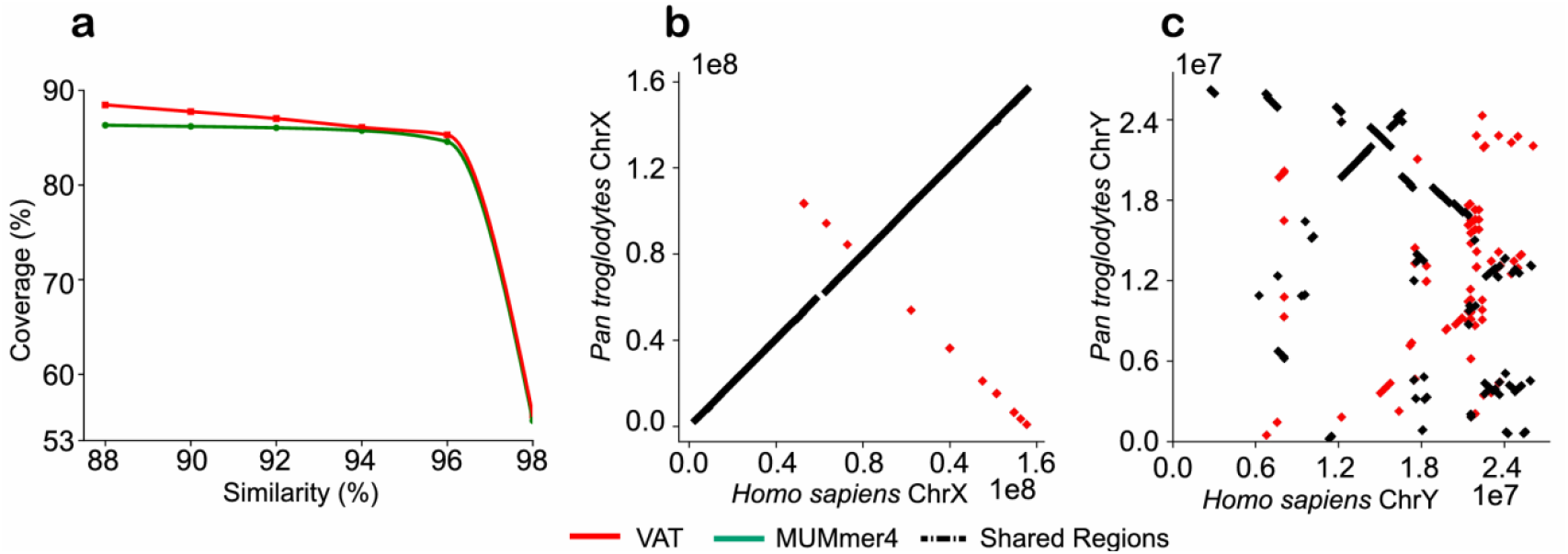
Performance of VAT and MUMmer4 on human–chimpanzee whole-genome alignment. (a) Alignment coverage across a range of sequence-identity thresholds for whole-genome alignments between *Homo sapiens* and *Pan troglodytes*, comparing VAT with MUMmer4. (b) Dot plot of alignments between human and chimpanzee chromosome X. (c) Dot plot of alignments between human and chimpanzee chromosome Y.

## 5. Discussion

We present VAT, a unified algorithmic framework for biological sequence alignment across scales and applications. Through simple run-time parameter tuning, VAT accommodates diverse alignment tasks across sequence alphabets (DNA and protein), lengths (from short-read mapping to whole-genome alignment), identities (from distant homology search to high-identity read mapping), structure (contiguous and split), and scales (from pairwise alignment to NT/NR-scale search). Despite this broad versatility, VAT achieves alignment quality and computational efficiency comparable to leading task-specific aligners. These results demonstrate the feasibility of combining high performance and generality within a single, unified algorithmic framework. Practically, VAT reduces the need to develop new aligners for emerging data types to the simpler task of parameter optimization or minor component refinement. As a multi-purpose aligner, VAT also streamlines the design of bioinformatic pipelines. For example, VAT enables the use of a single, unified human-genome index for both read mapping and whole-genome alignment between human and chimpanzee, without requiring task-specific indexing strategies, while in microbiome analyses, VAT can seamlessly integrate read mapping (16S-based taxonomic profiling), homology search (functional annotation of novel coding sequences), and whole-genome alignment (taxonomic assignment of assembled contigs).

