## Supplementary Table for "A general and extensible algorithmic framework to biological sequence alignment across scales and applications"

### 1. Supplementary Methods

#### 1.1. Multi-view table construction

VAT implements a unified multi-view seed index in which each row encodes a unique key string and each column stores positional entries contributed by a specific seeding scheme (e.g., $k$-mer, spaced seed/pattern, minimizer, and reduced-alphabet)[1-5]. All seed keys are compactly stored using a fixed-width binary representation: nucleotides are encoded using 2 bits (A, C, G, T/U), and amino acids are encoded using 5 bits. The 5-bit encoding accommodates 32 unique symbols, of which 20 represent standard amino acids and 10 represent reduced-alphabet groups used for sensitive protein homology search. Each key string is packed into a 64-bit integer. This corresponds to key string lengths of 32 nt or 12 aa, which provide sufficient discriminative power for the majority of DNA, RNA, and protein alignment tasks. The uniform 64-bit representation enables VAT to combine heterogeneous seeding strategies within the same index table without separate data structures or scheme-specific implementations.

During reference indexing, VAT performs a single-pass scan over the input database while simultaneously applying all supported seed types—exact $k$-mer, minimizer, spaced-pattern, and reduced-alphabet seeds. This unified traversal eliminates redundant I/O and ensures that all seed-derived positional information is captured concurrently. For exact $k$-mers, minimizers, and reduced-alphabet seeds, the key string is computed directly from the corresponding substring (native or alphabet-transformed). For spaced-pattern seeds, VAT extracts only residues at unmasked positions (designated as '1' in the pattern) to form the key string. The minimizer seeding scheme is defined by parameters $k$ and $h$: the minimizer is the lexicographically smallest $k$-mer within each window of length $h$. VAT fixes $k$ to the 64-bit key capacity (32 nt or 12 aa), while $h$ is varied across a user-defined range (32-42 for nt; 12-22 for aa). When $h = k$, the scheme reduces to exact $k$-mer matching. VAT stores minimizer-derived genomic coordinates for each specific $h$ value in dedicated columns of the multi-view index table. For spaced-pattern seeds, VAT supports 10 distinct mask patterns, each defined by its unique combination of masked ('0') and unmasked ('1') positions. Positional entries for each pattern are stored in separate columns. For reduced-alphabet seeding, VAT supports three distinct encoding schemes (e.g., Murphy-10[5], DIAMOND-11[6], and Thorne–Doolittle-10[7]), each mapping amino acids to reduced symbol classes prior to key computation. VAT records positional information for each reduced-alphabet scheme in its corresponding column of the multi-view table.

After positional information is generated for all seed types, VAT sorts the multi-view index table by the lexicographical order of the 64-bit key strings. We first apply a radix-based clustering approach[8] to partition key strings according to their 10-bit prefixes, yielding 1024 prefix buckets. This bucket count provides an effective balance between partition granularity, cache locality, and load balancing during parallel sorting. Within each prefix bucket, VAT performs an in-register SIMD bitonic sort to accelerate key ordering. Each table row is encoded in 128 bits: 64 bits encode the key string, and 64 bits store a pointer to positional records maintained across all seeding views. Using AVX-256 instructions, a 256-bit register holds two such rows simultaneously. The bitonic network operates on two registers at a time, enabling the sorting of four key strings (and their associated pointers) each time (see Figure 1 in the main article). This organization minimizes memory traffic and exploits data-level parallelism to achieve high-throughput key ordering during index construction.

Specifically, within each prefix bucket, VAT applies a fully in-register SIMD bitonic sorting pipeline operating on fixed-width key–pointer pairs. The sorter first partitions the bucket into vector-sized blocks and loads these blocks into AVX-256 registers. Each register holds two 128-bit seed records, enabling four key–pointer pairs to be processed per compare–exchange operation. The sorting process begins by forming local bitonic runs through sequences of in-register MIN/MAX comparisons. These comparisons reorder pairs of elements within each SIMD lane while maintaining pointer alignment. Inter-register compare–exchange steps then extend the bitonic structure across multiple registers, allowing runs to grow beyond the initial block boundaries. After each vertical comparison stage, a SIMD transpose is applied to rearrange elements so that logically adjacent keys become physically adjacent in SIMD lane order. This horizontal reorganization ensures proper alignment of elements for subsequent bitonic merging steps. Once bitonic runs of increasing size (e.g., 8, 16, 32, … up to the full bucket size) are constructed, the algorithm invokes a hierarchical set of SIMD merge networks—implemented using SHUFFLE, MIN, and MAX instructions—to merge sorted subsequences into progressively larger sorted segments. Iterating through all stages of the bitonic network yields a globally sorted ordering of key strings within each bucket, which are then concatenated to produce the final lexicographically ordered multi-view index.

After obtaining the fully sorted multi-view table, VAT computes the longest common prefix (LCP) between every adjacent pair of key strings to facilitate downstream query acceleration. For each key string $s_i$, VAT compares its 64-bit packed representation with that of its predecessor $s_{i-1}$ using a bit-parallel procedure. The two encoded keys are XORed to identify differing bit positions. The most significant set bit (MSB) in the XOR result marks the first mismatch between the two keys; its position is converted into a character-level prefix length based on the alphabet's bit width (2 bits for nucleotides, 5 bits for amino acids). This constant-time bitwise operation efficiently computes the LCP array across the entire index and enables VAT to skip redundant prefix regions during lookup, merge, and traversal operations.

In practice, VAT indexes the NR protein database (326 GB) in approximately 9.4 hours of wall-clock time using 32 threads, with a peak memory footprint of 14.6 GB. The resulting multi-view index occupies approximately 4.2 TB of disk space. Similarly, indexing the NT nucleotide database (1.3 TB) completes in 32.6 hours using 32 threads, requires 11.9 GB peak memory, and produces a final multi-view table of size 8.8 TB on disk.

#### 1.2. Seed matching and extension

VAT performs seed matching by simultaneously streaming the reference and query multi-view index tables and applying a modified merge-join algorithm[1,6,8]. This streaming design allows VAT to traverse both sorted tables in lockstep, efficiently identifying matching key strings across all seeding schemes without random memory access. Although VAT encodes fixed-length keys (32 nt for nucleotide seeds and 12 aa for protein seeds), it supports user-defined seed lengths shorter than the indexed key length. Seed lengths exceeding 32 nt or 12 aa are rarely useful in practical alignment scenarios, and the fixed 64-bit key representation provides sufficient resolution for typical DNA, RNA, and protein homology search tasks.

Specifically, to support seed lengths shorter than the fixed 32-nt or 12-aa indexed keys without regenerating the index, VAT performs an LCP-guided prefix search over its lexicographically sorted key strings. The LCP value between adjacent entries specifies the number of shared leading characters. For a user-specified seed length $l$, VAT locates the maximal contiguous block of keys whose pairwise minimum LCP is at least $l$; all keys in this block share an $l$-long prefix and jointly represent the complete set of occurrences of that shorter seed. VAT then retrieves the associated genomic or proteomic positions by taking the union of the positional lists from all keys within the LCP-defined block. Because each positional list is stored in sorted order, the union operation is performed via linear-time merging, ensuring efficient retrieval of all seed matches without reindexing.

After enumerating all candidate seeds, VAT applies a two-stage refinement strategy combining X-drop ungapped extension with SIMD-accelerated local alignment[9,10]. First, VAT performs an ungapped extension of each seed using the X-drop heuristic, which terminates extension when the running score falls more than X below the current maximum. This efficiently filters out weak or spurious matches. Only seeds whose ungapped extension scores exceed a predefined threshold proceed to gapped refinement. For gapped extension, VAT employs a striped Smith–Waterman algorithm in which match, mismatch, and gap scores are computed in parallel across SIMD lanes[9,10]. The striped layout ensures that data for multiple dynamic programming cells are simultaneously loaded and updated within AVX registers, substantially accelerating per-seed local alignment. Seeds producing high gapped-alignment scores are retained for downstream seed chaining, while low-scoring candidates are discarded.

#### 1.3. Chaining and gap-filling

VAT determines the optimal ordering of refined seeds using a dynamic-programming (DP) formulation[2,11]. Seed pairs are partially ordered by their query coordinates, and VAT models the cost of linking two seed pairs using a weighted linear scoring function. Specifically, to extend the existing seed chain ending at seed pair $j$ with a candidate seed pair $i$ ($i > j$), VAT employs a weighted linear scoring model that captures both alignment quality and positional consistency. The model incorporates $S_i$ (the matching score of seed $i$), $d_{i,q}$ (the query coordinate distance between seeds $j$ and $i$,), $d_{i,r}$ (the reference coordinate distance between seeds $j$ and $i$), and $p_i$ (a pattern-dependent bonus or penalty reflecting biologically meaningful sequence motifs). The DP recurrence maximizing the chain compatibility score $C$ is:

$$C[i] = \max_{i-j \le m} \{C[j] + w_1 S_i + w_2 d_{i,q} + w_3 |d_{i,r} - d_{i,q}| + w_4 p_i\}$$

The term $| d_{i,r} - d_{i,q} |$ approximates the implied gap size between seed pairs. For inter-chromosomal seed candidates, this value is set to $+\infty$; choosing $w_3 = 0$ thus permits inter-chromosomal chaining, which is required for applications such as CLASH (miRNA–mRNA chimeric junctions)[11,12]. The motif-dependent term $p_i$ (with tunable sequence pattern and the associated bonus/penalty scores) enables incorporation of biologically relevant signals, such as GT–AG splice sites or AAGCTT–GATC Hi-C ligation motifs[13], improving chain quality for RNA-seq and proximity-ligation data. The parameter $m$ limits the number of preceding seeds considered during DP. Larger $m$ values improve sensitivity and chain quality at the cost of increased runtime, whereas smaller $m$ values provide faster execution with reduced sensitivity.

After seed chaining, VAT performs gap filling between adjacent chained seeds using a SIMD-accelerated Needleman–Wunsch algorithm[14,15].

##### 1.4. Output format

VAT adopts the SAM format for reporting both short- and long-read alignments. Mapping quality (MAPQ) is computed using the same Phred-scaled formulation as BWA-MEM[16]. For homology search, VAT produces BLAST-like tabular outputs and computes E-values following the standard BLAST statistical model. For whole-genome alignment, VAT outputs MAF format and computes alignment score and percent identity as a similarity metric for evaluating correspondence between aligned synteny blocks.

### 2. VAT Parameter Setting for Different Alignment Applications

#### 2.1. All tunable parameters and their default values

- Seed length: (default: 14 for DNA; 8 for protein)
- Minimizer window size: (default: 0)
- Reduced alphabet: reduced amino acid alphabet (default: Murphy.10)
- Pattern number: number of spaced seed patterns (default: 1)
- Match: (default: +5 for DNA; BLOSUM62 for protein)
- Mismatch: (default: -3; BLOSUM62 for protein)
- Gap extension penalty: (default: -1)
- Gap open penalty: (default: −2)
- X-drop threshold ungapped (default: 18)
- X-drop threshold gapped: (default: 18)
- Min. chain length: minimum number of seeds required in a seed chain (default: 2)
- Seed dist. query: maximum distance allowed between two seeds in query (default: 150)
- Seed dist. reference: maximum distance allowed between two seeds in query (default: 150)
- Seed seq. pattern upstream: the sequence pattern expected from the upstream of the seeds and its associated bonus/penalty score (default: sequence = None; score = 0)
- Seed seq. pattern downstream: the sequence pattern expected from the downstream of the seeds and its associated bonus/penalty score (default: sequence = None; score = 0)
- Inter-chromosome chaining: (default: disallowed)
- Seed padding: successive seeds within same diagonal band skipped (10)
- E-value: maximum e-value to report alignments (default: 0.001)
- Report identity: minimum identity (%) to report an alignment (default: 0)
- Max. target sequences: maximum number of target sequences to report alignments (default: 25)

#### 2.2. Next-generation sequencing reads (NGS) mapping

##### 2.2.1. Contiguous (WGS, ChIP-seq, ATAC-seq, and 16S microbiome)

- Seed length: 15
- Minimizer window size: 5
- Match: +5
- Mismatch: −2

##### 2.2.2. RNA-seq

- Seed length: 14
- Minimizer window size: 5
- Match: +1
- Mismatch: -2
- X-drop threshold ungapped: 15
- X-drop threshold gapped: 15
- Seed dist. query: 1000
- Seed dist. reference: 200000
- Seed seq. pattern upstream: GT, +4
- Seed seq. pattern downstream: AG, +4

##### 2.2.3. circRNA

- Seed length: 14
- Minimizer window size: 2
- Match: +1
- Mismatch: -3
- Gap extension penalty: -2

- Gap open penalty: -5
- X-drop threshold ungapped: 15
- X-drop threshold gapped: 15
- Seed dist. query: 1000
- Seed dist. reference: 200000
- Seed seq. pattern upstream: AG, +4
- Seed seq. pattern downstream: GT, +4

###### 2.2.4. CLASH

- Seed length: 11
- Minimizer window size: 1
- Match: +5
- Mismatch: -4
- Gap extension penalty: -3
- Gap open penalty: -5
- X-drop threshold ungapped: 10
- X-drop threshold gapped: 10
- Inter-chromosome chaining: allowed

#### 2.2.5. Hi-C

- Seed length: 13
- Minimizer window size: 5
- Match: +1
- Mismatch: -3
- Gap extension penalty: -1
- Gap open penalty: -6
- Seed dist. query: 5000
- Seed dist. reference: 5000
- Seed seq. pattern upstream: AAGCTT, +4
- Seed seq. pattern downstream: GATC, +4
- Inter-chromosome chaining: allowed

##### 2.3. Third-generation sequencing read (TGS) mapping

###### 2.3.1. Contiguous (WGS, ChIP-seq, ATAC-seq, and 16S microbiome)

- Seed length: 15
- Minimizer window size: 10
- Mismatch: -4
- Match: +2
- Gap open penalty: -8

###### 2.3.2. RNA-seq

- Seed length: 15
- Minimizer window size: 5
- Match: +1
- Mismatch: -2
- Gap extension penalty: -1
- Gap open penalty: -2
- X-drop threshold ungapped: 20
- X-drop threshold gapped: 20
- Seed dist. query: 2000
- Seed dist. reference: 200000

- Seed seq. pattern upstream: GT, +4
- Seed seq. pattern downstream: AG, +4

#### 2.4. Homology search

##### 2.4.1. DNA

- Seed length: 15
- Minimizer window size: 3
- Match: +5
- Mismatch: -4
- Gap extension penalty: -3
- Gap open penalty: -5

##### 2.4.2. Protein (sensitive mode)

- Pattern number: 10
- Gap extension penalty: -1
- Gap open penalty: -11
- Seed padding: 16

##### 2.4.3. Protein (fast mode)

- Pattern number: 3
- Gap extension penalty: -1
- Gap open penalty: -11
- Seed padding threshold: 16

#### 2.5. Whole-genome alignment

- Seed length: 18
- Minimizer window size: 10
- Match: +1
- Mismatch: -5
- Gap extension penalty: -2
- Gap open penalty: -5
- X-drop threshold ungapped: 24
- X-drop threshold gapped: 24
- Seed padding threshold: 55

### 3. Benchmark Experiment Design

#### 3.1. NGS

##### 3.1.1. Contiguous

- Benchmark datasets: see Supplementary Table S1
- Ground-truth definition:
  - Read origin (simulated data only): recorded by the simulator Mason[17] during read synthesis.
- Benchmark metrics:
  - Overall mapping rate (simulated data only): proportion of reads successfully mapped to the reference (overlapping > 60% with ground-truth origin).
  - Accuracy: ratio of correctly aligned reads among all aligned reads.
  - Alignment speed: number of reads mapped per second.
  - Minimum alignment identity: lowest acceptable percent identity among aligned segments.
  - Minimum alignment length: shortest aligned region among reported segments.
  - Memory usage: peak RAM consumption (GB).

##### 3.1.2. Split

- Benchmark datasets: Supplementary Table S13
- Ground-truth definition:
  - Simulated data: read origins recorded by BEERS2[18] during synthesis.
  - RNA-seq: split reads on the same chromosome/orientation with ≤ 10 kbp distance whose junctions fully overlap annotated splice sites (Ensembl GRCh38[19]).
  - circRNA-seq: back-splicing junctions with reversed segment order and ≤ 200 kbp distance; verified against circAtlas 3.0 (~769k human circRNAs)[20].
  - CLASH: miRNA–mRNA duplexes where both arms overlap < 4 bp (Hyb "perfect" criterion); one arm on miRNA, the other on mRNA 3′ UTR[11,12,21,22].
  - Hi-C: split reads on the same chromosome/orientation with ≤ 200 kbp distance spanning annotated chromatin loops from Rao et al. (~10k loops)[13].
- Benchmark metrics:
  - Overall mapping rate: proportion of reads successfully mapped to ground truth.
  - Accuracy: ratio of correctly aligned reads among all aligned reads.
  - Alignment speed: reads mapped per second.
  - Minimum alignment identity: lowest acceptable percent identity among aligned segments.
  - Minimum alignment length: shortest aligned region among reported segments.
  - Memory usage (real data): peak RAM (GB).
  - Known proportion (split-signal quality): ratio of reads spanning annotated split events to all reads spanning split signals.

#### 3.2. TGS

##### 3.2.1. Contiguous

- Benchmark datasets: see Supplementary Table S25
- Ground-truth definition:
  - Read origin (simulated data only): recorded by the simulator Badread[23] during read synthesis.
- Benchmark metrics:
  - Overall mapping rate: proportion of reads successfully mapped to the reference (overlapping > 60% with ground-truth origin).
  - Accuracy: ratio of correctly aligned reads among all aligned reads.
  - Alignment speed: number of reads mapped per second.
  - Minimum alignment identity: lowest acceptable percent identity among aligned segments.
  - Minimum alignment length: shortest aligned region among reported segments.
  - Memory usage (real data): peak RAM consumption (GB).

##### 3.2.2. Split

- Benchmark datasets: see Supplementary Table S37
- Ground-truth definition:
  - Simulated data: read origins recorded by PBSIM[24] during synthesis.

- RNA-seq: split reads on the same chromosome/orientation with ≤ 10 kbp inter-segment distance, whose junctions fully overlap annotated splice sites (Ensembl GRCh38).

- Benchmark metrics:
  - Overall mapping rate: proportion of reads successfully mapped to the reference.
  - Known alignments (TGS simulated data): reads spanning multiple junctions with segments correctly matched to ground-truth loci.
  - Accuracy: ratio of correctly aligned reads among all aligned reads.
  - Alignment speed: number of reads mapped per second.
  - Minimum alignment identity: lowest acceptable percent identity among aligned segments.
  - Minimum alignment length: shortest aligned region among reported segments.
  - Memory usage (real data): peak RAM consumption (GB).
  - Known proportion (real data): reads with segments mapped to loci consistent with Ensembl GRCh38 annotation.

#### 3.3. Homology search

##### 3.3.1. Protein

- Benchmark datasets: target and query sequences derived from Pfam v37.0[25,26].
  - Sequences were randomly sampled with a cap per Pfam family to maintain diversity.
  - Selected sequences were split into two groups—query set and target database.
- Ground-truth definition:
  - Pfam-A curated families serve as the ground truth; two sequences are considered homologous if they belong to the same Pfam family.
  - Each Pfam family represents evolutionarily related proteins sharing conserved structural and functional domains.
- Benchmark metrics:
  - TP: query and hit from the same curated family (Pfam).
  - FP: hit from a different family.
  - FN: homologous family member not retrieved.
  - Sensitivity: completeness of retrieved homologues.

$$Sensitivity = \frac{TP}{TP + FN}$$

  - Precision: reliability of reported hits.

$$Precision = \frac{TP}{TP + FP}$$

  - F1 score: harmonic means of precision and sensitivity.

$$Precision = 2 \times \frac{Precision \times Sensitivity}{Precision + Sensitivity}$$

  - Alignment speed: number of queries aligned per second.
  - Memory usage: peak RAM consumption (GB).

##### 3.3.2. DNA

- Benchmark datasets: transcript sequences from the human reference genome used as the reference; query sets generated from known orthologous gene mappings between human and multiple species, including vertebrate and invertebrate model organisms.
- Ground-truth definition:
  - The Homologous Gene Database (HGD)[27] serves as the curated ground truth.
  - Each HGD family contains manually annotated orthologous and paralogous genes, representing validated homologous relationships across species.
- Benchmark metrics:
  - TP: query and hit from the same curated family (HGD).
  - FP: hit from a different family.
  - FN: homologous family member not retrieved.
  - Sensitivity: completeness of retrieved homologues.

$$Sensitivity = \frac{TP}{TP + FN}$$

- Precision: reliability of reported hits.

$$Precision = \frac{TP}{TP + FP}$$

- F1 score: harmonic means of precision and sensitivity.

$$Precision = 2 \times \frac{Precision \times Sensitivity}{Precision + Sensitivity}$$

- Alignment speed: number of queries aligned per second.
- Memory usage: peak RAM consumption (GB).

Supplementary Table S1 : Summary of contiguous next-generation sequencing and reference datasets used in this study, including simulated data and corresponding command-line parameters. The columns represent the dataset label (Label), sequence continuity status (SCS), data type, source organism (Organism), number of reads (# Reads), average read length (Len.), and accession numbers. *H. sapiens* refers to *Homo sapiens*, Gut MG refers to Gut Metagenome, *C. elegans* (*Caenorhabditis elegans*), *D. melan* (*Drosophila melanogaster*), *M. mus* (*Mus musculus*), *O. sativa* (*Oryza sativa*), and *A. thalia* (*Arabidopsis thaliana*). Specifically, NGS-Sim-DS1 is a simulated whole-genome sequencing dataset with an introduced error rate of 0.4%; NGS-Sim-DS2 is a simulated RNA-Seq dataset with an error rate of 0.6%.

| Label | SCS | Data type | Organism | # Reads | Len. | Accession | Simulation parameters |
|---|---|---|---|---|---|---|---|
| **NGS-Sim-DS1 (Mason2 v2.0.9)** | Contiguous | Simulation | *H. sapiens* | 5M | 150 | N/A | mason_simulator -ir GRCh38_genomic.fna -n 5000000 --illumina-read-length 150 -o hg38_reads.fa -oa hg38.reads.sam --illumina-prob-mismatch-scale 2.5 |
| **NGS-DS1** | Contiguous | WGS | *H. sapiens* | 513M | 151 | ERR1341796 | N/A |
| **NGS-DS2** | Contiguous | ChIP-Seq | *H. sapiens* | 13.8M | 150 | SRR32926125 | N/A |
| **NGS-DS3** | Contiguous | ATAC-seq | *H. sapiens* | 19.4M | 50 | SRR33004526 | N/A |
| **NGS-DS4** | Contiguous | Amplicon | Gut MG | 0.15M | 301 | SRR27677828 | N/A |
| **NGS-DS5** | Contiguous | WGS | *C. elegans* | 31.3M | 149 | SRR32699720 | N/A |
| **NGS-DS6** | Contiguous | WGS | *D. melan* | 8.0M | 151 | SRR32469661 | N/A |
| **NGS-DS7** | Contiguous | WGS | *M. mus* | 238.5K | 150 | SRR32563845 | N/A |
| **NGS-DS8** | Contiguous | WGS | *O. sativa* | 32.3M | 149 | ERR13765494 | N/A |
| **NGS-DS9** | Contiguous | WGS | *A. thalia* | 42.6M | 99 | SRR32329200 | N/A |

Supplementary Table S2: Summary of short-read contiguous aligners, their software versions, and the specific command-line parameters used for benchmarking in this study.

| Aligner | Version | Command |
|---|---|---|
| **BBMAP** | 35.85 | threads=16 |
| **BLASR** | 2012 | -nproc 16 -sam -out |
| **Bowtie2** | 2.5.4 | -L 15 -p 16 -U |
| **BWA MEM** | 0.7.17-r1198-dirty | -t 16 -k 15 |
| **HISAT2** | 2.2.0 | --pen-noncansplice 0 –all -p 16 --no-spliced-alignment |
| **Minimap2** | 2.30 (r1287) | -ax sr -k 15 -w 8 -t 16 |
| **SOAP2** | 2.21 | -p 16 |
| **STAR** | 2.7.11b | --genomeSAindexNbases 6 --runThreadN 16 |
| **segemehl** | 0.2.0-418 | -s -t 16 |
| **Tophat2** | 2.1.1 | -p 16 |
| **Subread** | 2.0.2 | -t 1 -T 16 -SAMoutput |
| **VAT** | 0.0.1 | -p 16 -wgs |

Supplementary Table S3: Benchmark of aligners on simulated contiguous NGS reads from *Homo sapiens*. Metrics include mapping rate (Map rate, %), accuracy (%), minimum alignment identity (Min identity, %), minimum alignment length (Min length), alignment speed (Speed, aligned reads per second), and peak memory usage (RAM, Gb).

| Aligner | Map rate | Accuracy | Min identity | Min length | Speed | RAM |
|---|---|---|---|---|---|---|
| **VAT** | 99.79 | 98.88 | 93.91 | 118 | 31112 | 14.12 |
| **BBMAP** | 95.51 | 98.53 | 87.89 | 51 | 3176 | 8.38 |
| **BLASR** | **100.00** | 97.07 | 84.02 | 40 | 583 | 29.88 |
| **Bowtie2** | 93.97 | 97.57 | 79.78 | 34 | 10800 | **3.68** |
| **BWA MEM** | 98.33 | 98.59 | 80.12 | 32 | 12721 | 11.03 |
| **HISAT2** | 97.01 | **99.05** | **94.94** | **120** | 39715 | 4.48 |
| **Minimap2** | 95.42 | 98.51 | 93.87 | 26 | 14599 | 11.32 |
| **SOAP2** | 58.10 | 98.11 | 94.77 | 118 | **42102** | 6.69 |
| **STAR** | 98.79 | 97.85 | 87.89 | 100 | 28193 | 34.21 |
| **segemehl** | 94.94 | 97.36 | 80.14 | 71 | 22353 | 32.13 |
| **subread** | 89.92 | 98.58 | 85.73 | 43 | 21818 | 7.96 |
| **Tophat2** | 94.49 | 98.59 | 90.92 | 110 | 2968 | 10.92 |

Supplementary Table S4: Benchmark of aligners on WGS contiguous NGS reads from *Homo sapiens*. Metrics include mapping rate (Map rate, %), minimum alignment identity (Min identity, %), minimum alignment length (Min Length), alignment speed (Speed, aligned reads per second), and peak memory usage (RAM, Gb).

| Aligner | Map rate | Min identity | Min length | Speed | RAM |
|---|---|---|---|---|---|
| **VAT** | **100.00** | 81.31 | 133 | **25222** | 15.01 |
| **BBMAP** | 78.23 | 80.94 | 100 | 9653 | 9.68 |
| **BLASR** | 99.82 | 80.92 | 38 | 593 | 21.04 |
| **Bowtie2** | 50.09 | 85.88 | 121 | 2575 | **3.81** |
| **BWA MEM** | **100.00** | 83.77 | 30 | 3856 | 20.79 |
| **HISAT2** | 89.94 | 79.98 | 128 | 22622 | 4.79 |
| **Minimap2** | **100.00** | 82.13 | 25 | 9000 | 16.94 |
| **SOAP2** | 13.22 | **88.23** | 120 | 8211 | 6.68 |
| **STAR** | **100.00** | 81.04 | 100 | 15426 | 30.77 |
| **segemehl** | 18.19 | 80.41 | 72 | 702 | 36.89 |
| **subread** | 27.64 | 81.33 | 15 | 6664 | 8.14 |
| **Tophat2** | 38.48 | 82.95 | 102 | 76 | 11.12 |

Supplementary Table S5: Benchmark of aligners on CHIP-seq contiguous NGS reads from *Homo sapiens*. Metrics include mapping rate (Map rate, %), minimum alignment identity (Min identity, %), minimum alignment length (Min length), alignment speed (Speed, aligned reads per second), and peak memory usage (RAM, Gb).

| Aligner | Map rate | Min identity | Min length | Speed | RAM |
|---|---|---|---|---|---|
| **VAT** | 98.98 | 83.33 | 113 | 25111 | 6.99 |
| **BBMAP** | 96.08 | 77.89 | 103 | 791 | 8.74 |
| **BLASR** | 98.96 | 79.78 | 40 | 359 | 29.98 |
| **Bowtie2** | 85.24 | 83.97 | 101 | 5427 | **3.69** |
| **BWA MEM** | **99.37** | 82.13 | 30 | 9119 | 9.27 |
| **HISAT2** | 78.02 | **85.32** | **124** | **25161** | 4.64 |
| **Minimap2** | 98.37 | 83.03 | 25 | 5927 | 17.08 |
| **SOAP2** | 68.92 | 81.77 | 84 | 10299 | 6.45 |
| **STAR** | 97.97 | 78.42 | 99 | 19216 | 31.18 |
| **segemehl** | 93.60 | 79.87 | 91 | 3520 | 17.88 |
| **subread** | 71.82 | 82.23 | 43 | 13296 | 21.04 |
| **Tophat2** | 84.33 | 80.98 | 108 | 271 | 4.14 |

Supplementary Table S6: Benchmark of aligners on ATAC-seq contiguous NGS reads from *Homo sapiens*. Metrics include mapping rate (Map rate, %), minimum alignment identity (Min identity, %), minimum alignment length (Min length), alignment speed (Speed, aligned reads per second), and peak memory usage (RAM, Gb).

| Aligner | Map rate | Min identity | Min length | Speed | RAM |
|---|---|---|---|---|---|
| **VAT** | 99.71 | 83.93 | 41 | 38250 | 16.12 |
| **BBMAP** | 97.79 | 79.02 | 38 | 8811 | 26.35 |
| **BLASR** | **100.00** | 79.77 | 37 | 449 | 29.82 |
| **Bowtie2** | 94.03 | 83.94 | **42** | 30323 | **3.49** |
| **BWA MEM** | 98.58 | 67.99 | 30 | 31797 | 6.94 |
| **HISAT2** | 88.10 | **85.60** | 41 | **44000** | 4.49 |
| **Minimap2** | 94.94 | 81.68 | 25 | 7840 | 16.92 |
| **SOAP2** | 86.68 | 80.34 | 36 | 22231 | 6.38 |
| **STAR** | 99.00 | 81.89 | 32 | 18000 | 29.70 |
| **segemehl** | 86.13 | 78.96 | 24 | 2385 | 30.88 |
| **subread** | 86.89 | 77.11 | 26 | 28065 | 6.83 |
| **Tophat2** | 87.62 | 81.03 | 28 | 292 | 7.18 |

Supplementary Table S7: Benchmark of aligners on 16S microbiome contiguous NGS reads. Metrics include mapping rate (Map rate, %), minimum alignment identity (Min identity, %), minimum alignment length (Min length), alignment speed (Speed, aligned reads per second), and peak memory usage (RAM, Gb).

| Aligner | Map rate | Min identity | Min length | Speed | RAM |
|---|---|---|---|---|---|
| **VAT** | 95.29 | 87.02 | **192** | **6313** | 2.43 |
| **BBMAP** | 90.29 | 85.42 | 184 | 564 | 14.89 |
| **BLASR** | **96.88** | 80.88 | 43 | 314 | 2.13 |
| **Bowtie2** | 93.01 | 87.93 | 114 | 6172 | **0.54** |
| **BWA MEM** | 95.48 | 80.67 | 40 | 3169 | 0.89 |
| **HISAT2** | 72.97 | 87.03 | 102 | 4845 | 1.87 |
| **Minimap2** | 95.03 | 80.94 | 50 | 3362 | 2.10 |
| **SOAP2** | 59.90 | **89.24** | 189 | 3539 | 1.57 |
| **STAR** | 48.93 | 84.43 | 170 | 426 | 28.88 |
| **segemehl** | 89.12 | 85.11 | 189 | 126 | 1.03 |
| **subread** | 24.08 | 88.32 | 28 | 984 | 6.72 |
| **Tophat2** | 71.14 | 84.77 | 184 | 208 | 1.31 |

Supplementary Table S8: Benchmark of aligners on contiguous NGS reads from *Caenorhabditis elegans*. Metrics include mapping rate (Map rate, %), minimum alignment identity (Min identity, %), minimum alignment length (Min length), alignment speed (Speed, aligned reads per second), and peak memory usage (RAM, Gb).

| Aligner | Map rate | Min identity | Min length | Speed | RAM |
|---|---|---|---|---|---|
| **VAT** | 99.81 | 89.22 | 119 | 79211 | 10.99 |
| **BBMAP** | 84.28 | 86.77 | 104 | 6965 | 13.31 |
| **BLASR** | 95.45 | 81.84 | 38 | 5302 | 27.88 |
| **Bowtie2** | 80.02 | 81.87 | 100 | 25813 | **3.97** |
| **BWA MEM** | 88.91 | 83.99 | 30 | 35564 | 10.88 |
| **HISAT2** | 76.85 | **90.68** | 105 | **85389** | 4.47 |
| **Minimap2** | 88.74 | 81.13 | 25 | 32867 | 11.89 |
| **SOAP2** | 85.22 | 88.32 | **121** | 9909 | 6.99 |
| **STAR** | **100.00** | 81.14 | 98 | 15873 | 28.13 |
| **segemehl** | 81.00 | 84.57 | 101 | 6694 | 32.41 |
| **subread** | 81.10 | 84.87 | 43 | 27613 | 9.27 |
| **Tophat2** | 72.03 | 89.13 | 110 | 800 | 12.03 |

Supplementary Table S9: Benchmark of aligners on contiguous NGS reads from *Drosophila melanogaster*. Metrics include mapping rate (Map rate, %), minimum alignment identity (Min identity, %), minimum alignment length (Min length), alignment speed (Speed, aligned reads per second), and peak memory usage (RAM, Gb).

| Aligner | Map rate | Min identity | Min length | Speed | RAM |
|---|---|---|---|---|---|
| **VAT** | 99.88 | **88.41** | 119 | **62133** | 13.41 |
| **BBMAP** | 81.35 | 81.01 | 85 | 4327 | 13.88 |
| **BLASR** | 99.41 | 80.94 | 52 | 5847 | 15.23 |
| **Bowtie2** | 68.85 | 83.76 | 111 | 20864 | 9.04 |
| **BWA MEM** | 98.08 | 84.88 | 30 | 30650 | 10.77 |
| **HISAT2** | 60.78 | 87.65 | **126** | 60780 | **4.93** |
| **Minimap2** | 98.07 | 88.14 | 25 | 28844 | 12.87 |
| **SOAP2** | 39.87 | 86.32 | 120 | 26580 | 6.92 |
| **STAR** | **100.00** | 82.79 | 100 | 25641 | 34.04 |
| **segemehl** | 66.03 | 83.67 | 109 | 2797 | 47.78 |
| **subread** | 78.80 | 80.13 | 43 | 26267 | 8.88 |
| **Tophat2** | 41.89 | 81.9 | 112 | 500 | 11.17 |

Supplementary Table S10: Benchmark of aligners on contiguous NGS reads from *Mus musculus*. Metrics include mapping rate (Map rate, %), minimum alignment identity (Min identity, %), minimum alignment length (Min length), alignment speed (Speed, aligned reads per second), and peak memory usage (RAM, Gb).

| Aligner | Map rate | Min identity | Min length | Speed | RAM |
|---|---|---|---|---|---|
| **VAT** | 99.73 | 89.67 | **39** | **19002** | 15.98 |
| **BBMAP** | 99.66 | 85.13 | **39** | 989 | 9.89 |
| **BLASR** | **100.00** | 83.77 | 31 | 132 | 20.67 |
| **Bowtie2** | 99.22 | 85.13 | **39** | 15755 | **3.89** |
| **BWA MEM** | 99.97 | 82.78 | 30 | 13228 | 22.77 |
| **HISAT2** | 97.86 | 86.98 | **39** | 16649 | 4.89 |
| **Minimap2** | 99.96 | 87.32 | 25 | 2454 | 16.81 |
| **SOAP2** | 96.02 | 87.41 | **39** | 6534 | 6.97 |
| **STAR** | **100.00** | 85.87 | **39** | 5954 | 31.04 |
| **segemehl** | **100.00** | 88.10 | **39** | 94 | 46.77 |
| **subread** | 69.50 | **90.90** | **39** | 10346 | 8.50 |
| **Tophat2** | 81.04 | 88.03 | **39** | 193 | 9.83 |

Supplementary Table S11: Benchmark of aligners on contiguous NGS reads from *Oryza sativa*. Metrics include mapping rate (Map rate, %), minimum alignment identity (Min identity, %), minimum alignment length (Min length), alignment speed (Speed, aligned reads per second), and peak memory usage (RAM, Gb).

| Aligner | Map rate | Min identity | Min length | Speed | RAM |
|---|---|---|---|---|---|
| **VAT** | 99.87 | **88.41** | **31** | 45800 | 15.89 |
| **BBMAP** | 96.96 | 83.67 | 30 | 4998 | 7.87 |
| **BLASR** | **100.00** | 84.03 | 29 | 1869 | 25.13 |
| **Bowtie2** | 94.38 | 83.89 | **31** | 21450 | **4.09** |
| **BWA MEM** | 98.54 | 86.60 | 30 | 19708 | 12.88 |
| **HISAT2** | 88.62 | 87.59 | 29 | **49233** | 4.98 |
| **Minimap2** | 98.60 | 83.87 | 25 | 17927 | 15.77 |
| **SOAP2** | 65.92 | 88.23 | 29 | 28661 | 6.60 |
| **STAR** | **100.00** | 80.16 | 24 | 41667 | 31.79 |
| **segemehl** | 92.89 | 82.98 | 28 | 3676 | 45.16 |
| **subread** | 66.50 | 80.76 | 22 | 22931 | 8.31 |
| **Tophat2** | 89.84 | 86.41 | 29 | 1800 | 10.47 |

Supplementary Table S12: Benchmark of aligners on contiguous NGS reads from *Arabidopsis thaliana.* Metrics include mapping rate (Map rate, %), minimum alignment identity (Min identity, %), minimum alignment length (Min length), alignment speed (Speed, aligned reads per second), and peak memory usage (RAM, Gb).

| Aligner | Map rate | Min identity | Min length | Speed | RAM |
|---|---|---|---|---|---|
| **VAT** | 99.89 | 88.91 | **37** | 81923 | 16.08 |
| **BBMAP** | 98.40 | 86.57 | 29 | 8708 | 9.10 |
| **BLASR** | **100.00** | 83.78 | 29 | 3356 | 20.78 |
| **Bowtie2** | 97.66 | 86.13 | 35 | 42461 | 4.79 |
| **BWA MEM** | 98.97 | 82.89 | 30 | 44986 | 10.48 |
| **HISAT2** | 92.94 | **89.91** | 35 | **92940** | **4.68** |
| **Minimap2** | 98.90 | 84.03 | 25 | 39560 | 13.58 |
| **SOAP2** | 85.75 | 86.10 | 30 | 50441 | 6.80 |
| **STAR** | **100.00** | 87.89 | 26 | 32258 | 29.33 |
| **segemehl** | 91.91 | 80.13 | 31 | 2621 | 33.79 |
| **subread** | 72.69 | 80.67 | 24 | 29080 | 9.17 |
| **Tophat2** | 88.78 | 87.41 | 31 | 1271 | 9.98 |

Supplementary Table S13: Summary of split next-generation sequencing and reference datasets used in this study, including simulated data and corresponding command-line parameters. The columns represent the dataset label (Label), sequence continuity status (SCS), data type, source organism (Organism), number of reads (# Reads), average read length (Len.), and accession numbers. *H. sapiens* refers to *Homo sapiens*, Gut MG refers to Gut Metagenome, *C. elegans* (*Caenorhabditis elegans*), *D. melan* (*Drosophila melanogaster*), *M. mus* (*Mus musculus*), *O. sativa* (*Oryza sativa*), and *A. thalia* (*Arabidopsis thaliana*). Specifically, NGS-Sim-DS1 is a simulated whole-genome sequencing dataset with an introduced error rate of 0.4%; NGS-Sim-DS2 is a simulated RNA-Seq dataset with an error rate of 0.6%.

| Label | SCS | Data type | Organism | # Reads | Len. | Accession | Simulation parameters |
|---|---|---|---|---|---|---|---|
| **NGS-Sim-DS2 (BEERS2)** | Split | Simulation | *H. sapiens* | 10M | 100 | N/A | perl reads_simulator.pl 10000000 T1_PE100 -readlength 100 -fraglength 100,250,500 -error 0.0061 -subfreq 0 -indelfreq 0 -mastercfgdir master_cfg -outdir T1_low |
| **NGS-DS10** | Split | RNA-Seq | *H. sapiens* | 108M | 101 | SRR534301 | N/A |
| **NGS-DS11** | Split | circRNA | *H. sapiens* | 13.3M | 101 | SRR1636985 | N/A |
| **NGS-DS12** | Split | CLASH | *H. sapiens* | 52.7M | 55 | SRR959751 | N/A |
| **NGS-DS13** | Split | Hi-C | *H. sapiens* | 1.52M | 95 | SRR1658825 | N/A |
| **NGS-DS14** | Split | RNA-Seq | *C. elegans* | 3.6M | 130 | SRR31943890 | N/A |
| **NGS-DS15** | Split | RNA-Seq | *D. melan* | 19.8M | 150 | SRR30712194 | N/A |
| **NGS-DS16** | Split | RNA-Seq | *M. mus* | 25.5M | 151 | SRR32754573 | N/A |
| **NGS-DS17** | Split | RNA-Seq | *O. sativa* | 19.7M | 150 | SRR32695903 | N/A |
| **NGS-DS18** | Split | RNA-Seq | *A. thalia* | 14.1M | 126 | SRR32771535 | N/A |

Supplementary Table S14: Summary of short-read split aligners, their software versions, and the specific command-line parameters used for benchmarking in this study.

| Aligner | Version | Command |
|---|---|---|
| **BBMAP** | 35.85 | threads=16 |
| **BLASR** | 2012 | -nproc 16 -sam -out |
| **Bowtie2** | 2.5.4 | -L 15 -p 16 -U |
| **BWA MEM** | 0.7.17-r1198-dirty | -t 16 -k 15 -k 4 -D 20 -R 3 -N 0 --score-min L,18,0 --end-to-end --rdg 5,1 --rfg 5,1 |
| **HISAT2** | 2.2.0 | -p 16 |
| **Minimap2** | 2.30 (r1287) | -k 15 -ax splice:hq -t 16 |
| **SOAP2** | 2.21 | -p 16 |
| **STAR** | 2.7.11b | --genomeSAindexNbases 6 --runThreadN 16 |
| **segemehl** | 0.2.0-418 | -s -t 16 |
| **Tophat2** | 2.1.1 | -p 16 |
| **Subread** | 2.0.2 | subjunc -t 0 -T 16 -SAMoutput |
| **VAT** | 0.0.1 | -p 16 -splice --short |

Supplementary Table S15: Benchmark of aligners on simulated split NGS reads from *Homo sapiens*. Metrics include mapping rate (Map rate, %), proportions of contiguous alignments (Contiguous, %), proportions of split alignments (Split, %), accuracy (%), proportion of known split events (Known, %), alignment speed (Speed, aligned reads per second), and peak memory usage (RAM, Gb).

| Aligner | Map rate | Contiguous | Split | Accuracy | Known | Speed | RAM |
|---|---|---|---|---|---|---|---|
| **VAT** | **98.70** | 83.98 | **14.72** | 97.27 | **87.96** | 62917 | 17.92 |
| **BBMAP** | 96.18 | 82.41 | 13.77 | 93.85 | 67.88 | 7433 | 13.31 |
| **BLASR** | 95.00 | 83.69 | 11.31 | 88.76 | 28.43 | 3248 | 29.78 |
| **Bowtie2** | 96.96 | **85.77** | 11.19 | 89.59 | 28.03 | 16876 | **3.96** |
| **BWA MEM** | 93.96 | 80.19 | 13.77 | 90.08 | 45.77 | 17334 | 8.85 |
| **HISAT2** | 98.62 | 84.33 | 14.29 | **97.45** | 86.82 | **83482** | 5.13 |
| **Minimap2** | 94.83 | 80.27 | 14.56 | 93.56 | 69.91 | 25295 | 22.94 |
| **SOAP2** | 86.03 | 75.62 | 10.41 | 89.08 | 23.17 | 42949 | 7.33 |
| **STAR** | 97.92 | 83.20 | **14.72** | 95.01 | 80.11 | 74992 | 33.97 |
| **segemehl** | 95.03 | 80.81 | 14.22 | 93.88 | 75.58 | 4602 | 34.67 |
| **subread** | 95.25 | 80.77 | 14.48 | 94.64 | 74.77 | 47136 | 9.28 |
| **Tophat2** | 96.39 | 83.58 | 12.81 | 94.85 | 70.62 | 2988 | 11.10 |

Supplementary Table S16: Benchmark of aligners on real circRNA split NGS reads from *Homo sapiens.* Metrics include mapping rate (Map rate, %), proportions of contiguous alignments (Contiguous, %), proportions of split alignments (Split, %), proportion of known split events (Known, %), alignment speed (Speed, aligned reads per second), and peak memory usage (RAM, Gb).

| Aligner | Map rate | Contiguous | Split | Known | Speed | RAM |
|---|---|---|---|---|---|---|
| **VAT** | **97.98** | 91.07 | **6.91** | **22.11** | 64062 | 18.11 |
| **BBMAP** | 88.78 | 84.89 | 3.89 | 14.33 | 11646 | 7.89 |
| **BLASR** | 95.81 | **93.81** | 2.00 | 15.71 | 7295 | 25.12 |
| **Bowtie2** | 96.99 | 91.91 | 5.08 | 19.88 | 20182 | **3.78** |
| **BWA MEM** | 95.77 | 89.80 | 5.97 | 20.34 | 27624 | 11.23 |
| **HISAT2** | 48.96 | 44.07 | 4.89 | 13.26 | 28968 | 4.84 |
| **Minimap2** | 67.00 | 64.03 | 2.97 | 18.94 | 34348 | 19.01 |
| **SOAP2** | 34.80 | 33.84 | 0.96 | 12.81 | 22008 | 7.13 |
| **STAR** | 90.03 | 87.05 | 2.98 | 16.44 | **64610** | 33.88 |
| **segemehl** | 92.04 | 85.91 | 6.13 | 19.24 | 10259 | 51.41 |
| **subread** | 88.50 | 83.47 | 5.03 | 18.46 | 12955 | 8.89 |
| **Tophat2** | 80.98 | 77.11 | 3.87 | 17.78 | 4569 | 10.15 |

Supplementary Table S17: Benchmark of aligners on real RNA-seq split NGS reads from *Homo sapiens*. Metrics include mapping rate (Map rate, %), proportions of contiguous alignments (Contiguous, %), proportions of split alignments (Split, %), proportion of known split events (Known, %), alignment speed (Speed, aligned reads per second), and peak memory usage (RAM, Gb).

| Aligner | Map rate | Contiguous | Split | Known | Speed | RAM |
|---|---|---|---|---|---|---|
| **VAT** | **99.33** | 62.81 | 36.52 | **94.27** | **21619** | 16.89 |
| **BBMAP** | 95.70 | 70.32 | 25.38 | 67.60 | 2344 | 11.88 |
| **BLASR** | 95.77 | **83.48** | 12.29 | 16.19 | 1361 | 26.93 |
| **Bowtie2** | 80.39 | 53.48 | 26.91 | 68.70 | 11551 | **3.89** |
| **BWA MEM** | 96.55 | 71.89 | 24.66 | 31.65 | 18182 | 8.95 |
| **HISAT2** | 90.31 | 56.84 | 33.47 | 93.01 | 19867 | 5.47 |
| **Minimap2** | 98.01 | 64.99 | 33.02 | 73.00 | 6250 | 19.89 |
| **SOAP2** | 57.61 | 38.18 | 19.43 | 0.11 | 9344 | 7.27 |
| **STAR** | 99.24 | 62.01 | **37.23** | 91.42 | 16667 | 34.03 |
| **segemehl** | 93.32 | 62.11 | 31.21 | 69.62 | 3084 | 37.91 |
| **subread** | 90.26 | 56.79 | 33.47 | 80.82 | 14095 | 9.13 |
| **Tophat2** | 84.22 | 55.52 | 28.70 | 74.18 | 1149 | 10.88 |

Supplementary Table S18: Benchmark of aligners on real CLASH split NGS reads from *Homo sapiens*. Metrics include mapping rate (Map rate, %), proportions of contiguous alignments (Contiguous, %), proportions of split alignments (Split, %), proportion of known split events (Known, %), alignment speed (Speed, aligned reads per second), and peak memory usage (RAM, Gb).

| Aligner | Map rate | Contiguous | Split | Known | Speed | RAM |
|---|---|---|---|---|---|---|
| **VAT** | 97.17 | 31.28 | **65.89** | **95.35** | **114514** | 8.91 |
| **BBMAP** | 74.18 | **74.18** | 0.00 | 0.00 | 25218 | 7.88 |
| **BLASR** | 65.21 | 65.21 | 0.00 | 0.00 | 7367 | 18.78 |
| **Bowtie2** | **98.15** | 39.91 | 58.24 | 95.31 | 46463 | 4.03 |
| **BWA MEM** | 96.95 | 41.04 | 55.91 | 94.22 | 9704 | 9.89 |
| **HISAT2** | 25.87 | 25.87 | 0.00 | 0.00 | 36593 | **3.21** |
| **Minimap2** | 90.85 | 35.89 | 54.96 | 90.42 | 79749 | 6.33 |
| **SOAP2** | 22.11 | 22.11 | 0.00 | 0.00 | 33668 | 7.19 |
| **STAR** | 55.50 | 49.93 | 5.57 | 25.00 | 7912 | 31.88 |
| **segemehl** | 64.08 | 63.50 | 0.58 | 17.11 | 7245 | 36.89 |
| **subread** | 68.01 | 55.99 | 12.02 | 25.41 | 34147 | 8.87 |
| **Tophat2** | 47.89 | 47.89 | 0.00 | 0.00 | 6601 | 7.98 |

Supplementary Table S19: Benchmark of aligners on real Hi-C split NGS reads from *Homo sapiens*. Metrics include mapping rate (Map rate, %), proportions of contiguous alignments (Contiguous, %), proportions of split alignments (Split, %), proportion of known split events (Known, %), alignment speed (Speed, aligned reads per second), and peak memory usage (RAM, Gb).

| Aligner | Map rate | Contiguous | Split | Known | Speed | RAM |
|---|---|---|---|---|---|---|
| **VAT** | **99.13** | 95.02 | 4.11 | **32.33** | 59955 | 16.33 |
| **BBMAP** | 91.26 | 88.37 | 2.89 | 20.63 | 3599 | 15.93 |
| **BLASR** | 98.10 | 95.17 | 2.93 | 22.79 | 1056 | 10.91 |
| **Bowtie2** | 98.48 | **96.11** | 2.37 | 30.36 | 4198 | 3.89 |
| **BWA MEM** | 98.76 | 95.89 | 2.87 | 31.96 | 21931 | 10.96 |
| **HISAT2** | 73.00 | 69.96 | 3.62 | 15.34 | 75924 | **2.66** |
| **Minimap2** | 98.01 | 95.88 | 2.13 | 32.25 | 17907 | 15.89 |
| **SOAP2** | 60.45 | 58.46 | 1.99 | 3.64 | 18708 | 7.78 |
| **STAR** | 98.08 | 91.97 | 6.11 | 18.22 | **82365** | 31.32 |
| **segemehl** | 89.82 | 83.43 | **6.39** | 26.94 | 28473 | 41.45 |
| **subread** | 80.43 | 76.40 | 4.03 | 25.90 | 24742 | 98.78 |
| **Tophat2** | 74.70 | 71.72 | 2.98 | 3.82 | 645 | 10.87 |

Supplementary Table S20: Benchmark of aligners on real RNA-seq split NGS reads from *Caenorhabditis elegans*. Metrics include mapping rate (Map rate, %), proportions of contiguous alignments (Contiguous, %), proportions of split alignments (Split, %), proportion of known split events (Known, %), alignment speed (Speed, aligned reads per second), and peak memory usage (RAM, Gb).

| Aligner | Map rate | Contiguous | Split | Known | Speed | RAM |
|---|---|---|---|---|---|---|
| **VAT** | **99.29** | 66.98 | 32.31 | **78.20** | 32097 | 12.94 |
| **BBMAP** | 97.22 | 77.03 | 20.19 | 50.63 | 6107 | 16.33 |
| **BLASR** | 96.98 | 61.87 | **35.11** | 51.01 | 1366 | 26.89 |
| **Bowtie2** | 87.01 | 75.89 | 11.12 | 12.51 | 20810 | **3.92** |
| **BWA MEM** | 99.03 | **86.14** | 12.89 | 19.97 | 24808 | 10.83 |
| **HISAT2** | 97.99 | 68.96 | 29.03 | 73.13 | **40829** | 4.88 |
| **Minimap2** | 96.98 | 79.09 | 17.89 | 38.51 | 20606 | 13.78 |
| **SOAP2** | 76.40 | 63.59 | 12.81 | 2.60 | 19110 | 8.82 |
| **STAR** | 22.10 | 19.02 | 3.08 | 2.81 | 7419 | 23.79 |
| **segemehl** | 93.99 | 70.95 | 23.04 | 53.13 | 5193 | 39.14 |
| **subread** | 72.31 | 51.03 | 21.28 | 51.75 | 5975 | 9.20 |
| **Tophat2** | 87.09 | 66.95 | 20.14 | 47.03 | 2433 | 10.37 |

Supplementary Table S21: Benchmark of aligners on real RNA-seq split NGS reads from *Drosophila melanogaster*. Metrics include mapping rate (Map rate, %), proportions of contiguous alignments (Contiguous, %), proportions of split alignments (Split, %), proportion of known split events (Known, %), alignment speed (Speed, aligned reads per second), and peak memory usage (RAM, Gb).

| Aligner | Map rate | Contiguous | Split | Known | Speed | RAM |
|---|---|---|---|---|---|---|
| **VAT** | **99.18** | 68.29 | **30.89** | **90.30** | **58824** | 15.87 |
| **BBMAP** | 89.99 | 61.76 | 28.23 | 75.02 | 5150 | 14.42 |
| **BLASR** | 93.02 | 79.98 | 13.04 | 18.21 | 1550 | 23.99 |
| **Bowtie2** | 88.09 | 75.20 | 12.89 | 14.29 | 25114 | **3.75** |
| **BWA MEM** | 92.99 | **80.05** | 12.94 | 32.50 | 19496 | 11.80 |
| **HISAT2** | 68.98 | 46.10 | 22.88 | 63.75 | 9200 | 4.74 |
| **Minimap2** | 88.95 | 59.91 | 29.04 | 79.86 | 13455 | 15.89 |
| **SOAP2** | 58.93 | 49.12 | 9.81 | 3.71 | 34247 | 8.30 |
| **STAR** | 98.68 | 67.91 | 30.77 | 90.04 | 16667 | 30.11 |
| **segemehl** | 96.97 | 74.01 | 22.96 | 63.75 | 3904 | 33.76 |
| **subread** | 68.82 | 41.72 | 27.10 | 77.14 | 18421 | 8.20 |
| **Tophat2** | 95.97 | 67.76 | 28.21 | 81.71 | 1811 | 10.94 |

Supplementary Table S22: Benchmark of aligners on real RNA-seq split NGS reads from *Mus musculus*. Metrics include mapping rate (Map rate, %), proportions of contiguous alignments (Contiguous, %), proportions of split alignments (Split, %), proportion of known split events (Known, %), alignment speed (Speed, aligned reads per second), and peak memory usage (RAM, Gb).

| Aligner | Map rate | Contiguous | Split | Known | Speed | RAM |
|---|---|---|---|---|---|---|
| **VAT** | **99.22** | 52.94 | **46.28** | **92.05** | 25641 | 17.20 |
| **BBMAP** | 97.90 | 60.01 | 37.89 | 72.15 | 4055 | 13.13 |
| **BLASR** | 98.98 | **72.19** | 26.79 | 24.59 | 317 | 28.27 |
| **Bowtie2** | 71.83 | 53.79 | 18.04 | 14.27 | 13646 | **4.02** |
| **BWA MEM** | 98.99 | 70.86 | 28.13 | 40.39 | 15348 | 8.89 |
| **HISAT2** | 90.87 | 46.99 | 43.88 | 84.88 | **37829** | 4.91 |
| **Minimap2** | 98.91 | 56.87 | 42.04 | 78.78 | 6599 | 17.93 |
| **SOAP2** | 51.89 | 41.01 | 10.88 | 1.90 | 11788 | 7.34 |
| **STAR** | 96.95 | 59.78 | 37.17 | 77.68 | 33333 | 34.21 |
| **segemehl** | 71.94 | 48.88 | 23.06 | 41.95 | 1941 | 40.00 |
| **subread** | 84.95 | 50.78 | 34.17 | 69.56 | 18620 | 8.98 |
| **Tophat2** | 80.76 | 54.99 | 25.77 | 49.37 | 970 | 10.80 |

Supplementary Table S23: Benchmark of aligners on real RNA-seq split NGS reads from *Oryza sativa*. Metrics include mapping rate (Map rate, %), proportions of contiguous alignments (Contiguous, %), proportions of split alignments (Split, %), proportion of known split events (Known, %), alignment speed (Speed, aligned reads per second), and peak memory usage (RAM, Gb).

| Aligner | Map rate | Contiguous | Split | Known | Speed | RAM |
|---|---|---|---|---|---|---|
| **VAT** | 99.21 | 65.94 | 33.27 | **91.12** | 66667 | 14.21 |
| **BBMAP** | 96.89 | 67.16 | 29.73 | 76.64 | 7017 | 10.85 |
| **BLASR** | 95.88 | **73.87** | 22.01 | 36.77 | 1362 | 26.78 |
| **Bowtie2** | 73.89 | 58.01 | 15.88 | 23.93 | 20960 | **3.82** |
| **BWA MEM** | 98.38 | 72.02 | 26.36 | 52.64 | 39612 | 9.89 |
| **HISAT2** | 92.97 | 66.94 | 26.03 | 66.03 | **85255** | 4.85 |
| **Minimap2** | 97.77 | 63.81 | 33.96 | 85.16 | 28888 | 16.12 |
| **SOAP2** | 58.85 | 45.97 | 12.88 | 4.35 | 36888 | 7.47 |
| **STAR** | **99.35** | 65.05 | **34.30** | 87.09 | 76923 | 31.86 |
| **segemehl** | 71.09 | 60.96 | 10.13 | 15.48 | 4329 | 42.13 |
| **subread** | 95.07 | 67.06 | 28.01 | 73.38 | 25158 | 9.30 |
| **Tophat2** | 88.78 | 60.88 | 27.90 | 68.12 | 2469 | 10.96 |

Supplementary Table S24: Benchmark of aligners on real RNA-seq split NGS reads from *Arabidopsis thaliana*. Metrics include mapping rate (Map rate, %), proportions of contiguous alignments (Contiguous, %), proportions of split alignments (Split, %), proportion of known split events (Known, %), alignment speed (Speed, aligned reads per second), and peak memory usage (RAM, Gb).

| Aligner | Map rate | Contiguous | Split | Known | Speed | RAM |
|---|---|---|---|---|---|---|
| **VAT** | **99.52** | 59.03 | **40.49** | **91.04** | 11111 | 13.11 |
| **BBMAP** | 97.79 | 61.01 | 36.78 | 81.78 | 859 | 9.88 |
| **BLASR** | 95.77 | 74.05 | 21.72 | 22.05 | 1053 | 24.23 |
| **Bowtie2** | 73.05 | 60.87 | 12.18 | 12.97 | 2459 | **3.98** |
| **BWA MEM** | 97.78 | **77.02** | 20.76 | 36.48 | 5445 | 12.78 |
| **HISAT2** | 95.08 | 56.98 | 38.10 | 83.24 | **14909** | 4.87 |
| **Minimap2** | 97.97 | 64.18 | 33.79 | 70.54 | 8099 | 15.21 |
| **SOAP2** | 59.17 | 47.99 | 11.18 | 2.81 | 3094 | 8.10 |
| **STAR** | 99.37 | 60.97 | 38.40 | 82.70 | 11236 | 30.31 |
| **segemehl** | 94.89 | 61.78 | 33.11 | 71.21 | 1836 | 40.89 |
| **subread** | 96.83 | 60.01 | 36.82 | 87.02 | 2288 | 8.90 |
| **Tophat2** | 91.98 | 54.89 | 37.09 | 80.86 | 618 | 10.93 |

Supplementary Table S25: Summary of the contiguous third-generation sequencing (TGS) datasets and simulation commands used in this study. The columns represent the dataset label (Label), sequence continuity status (SCS), data type, source organism (Organism), number of reads (# Reads), average read (Len.), and accession numbers. *H. sapiens* refers to *Homo sapiens*, Gut MG refers to Gut Metagenome, *C. elegans* (*Caenorhabditis elegans*), *D. melan* (*Drosophila melanogaster*), *M. mus* (*Mus musculus*), *O. sativa* (*Oryza sativa*), and *A. thalia* (*Arabidopsis thaliana*). Specifically, TGS-Sim-DS1 is a simulated whole-genome sequencing dataset with an error rate of 12%; TGS-Sim-DS2 is a simulated RNA-Seq dataset with an introduced error rate of 15%. TGS-Sim-DS3 is a simulated Hi-C dataset with an error rate of 15%.

| Label | SCS | Data type | Organism | # Reads | Len. | Accession | Simulation parameters |
|---|---|---|---|---|---|---|---|
| **TGS-Sim-DS1 (Badread)** | Contiguous | Simulation | *H. sapiens* | 0.5M | 2500 | N/A | badread simulate --reference GRCh38_genomic.fna --quantity 1x --error_model random --qscore_model ideal --glitches 0,0,0 --junk_reads 0 --random_reads 0 --chimeras 0 --identity 30,3 --length 20000,1000 --start_adapter_seq "" --end_adapter_seq "" |
| **TGS-DS1** | Contiguous | WGS | *H. sapiens* | 0.67M | ~10k | ERR2184700 | N/A |
| **TGS-DS2** | Contiguous | ChIP-Seq | *H. sapiens* | 0.88M | 4563 | SRR27030338 | N/A |
| **TGS-DS3** | Contiguous | ATAC-seq | *H. sapiens* | 31K | 4328 | SRR17667631 | N/A |
| **TGS-DS4** | Contiguous | Amplicon | Gut MG | 49K | 766 | SRR32253554 | N/A |
| **TGS-DS6** | Contiguous | WGS | *C. elegans* | 1.1M | 7063 | SRR32300787 | N/A |
| **TGS-DS7** | Contiguous | WGS | *D. melan* | 876.0K | 8610 | SRR31826012 | N/A |
| **TGS-DS8** | Contiguous | WGS | *M. mus* | 3.2M | 7621 | ERR12143047 | N/A |
| **TGS-DS9** | Contiguous | WGS | *O. sativa* | 496.2K | 16329 | ERR13336064 | N/A |
| **TGS-DS10** | Contiguous | WGS | *A. thalia* | 988.8K | 583 | ERR14129311 | N/A |

Supplementary Table S26: Summary of long-read contiguous aligners, their software versions, and the specific command-line parameters used for benchmarking in this study.

| Aligner | Version | Command |
|---|---|---|
| **BLASR** | 2012 | -nproc 16 -sam -bestn 10 |
| **GMAP** | 6/24/2024 | -f samse -t 16 |
| **GraphMap2** | 0.6.5 | -t 16 |
| **Minimap2** | 2.30 (r1287) | -ax map-ont -k 15 -t 16 |
| **ngmlr** | 0.2.7 | -x ont -t 16 |
| **STARlong** | 2.7.11b | --runThreadN 16 \<br>--seedSearchStartLmax 14 \<br>--seedPerReadNmax 100000 \<br>--seedPerWindowNmax 100 \<br>--winAnchorMultimapNmax 200 \<br>--outFilterMultimapNmax 100000 \<br>--outFilterMismatchNmax 100000 \<br>--outFilterScoreMin 0 --outFilterScoreMinOverLread 0 \<br>--outFilterMatchNmin 0 --outFilterMatchNminOverLread 0 \<br>--alignIntronMin 20 --alignIntronMax 1000000 \<br>--alignSJoverhangMin 8 --alignSJDBoverhangMin 1 \<br>--alignMatesGapMax 1000000 \<br>--clip3pAfterAdapterNbases 1 --clip5pNbases 0 \<br>--chimOutType WithinBAM SoftClip \<br>--outSAMtype SAM \<br>--limitBAMsortRAM 12000000000 |
| **VAT** | 0.0.1 | -wgs --long -p 16 |

Supplementary Table S27: Benchmark of aligners on simulated contiguous TGS reads from *Homo sapiens*. Metrics include mapping rate (Map rate, %), accuracy (%), minimum alignment identity (Min identity, %), minimum alignment length (Min length), alignment speed (Speed, aligned reads per second), and peak memory usage (RAM, Gb).

| Aligner | Map rate | Accuracy | Min identity | Min length | Speed | RAM |
|---|---|---|---|---|---|---|
| **VAT** | **96.81** | **98.60** | **91.13** | 259 | **57** | 23.43 |
| **BLASR** | 96.03 | 97.52 | 66.89 | 198 | 2 | 23.78 |
| **GMAP** | 94.29 | 98.06 | 89.87 | 248 | 3 | 13.31 |
| **GraphMap2** | 95.89 | 98.55 | 89.04 | 255 | 6 | 80.89 |
| **ngmlr** | 93.94 | 98.48 | 90.94 | 257 | 4 | **12.13** |
| **Minimap2** | 96.27 | 98.47 | 85.10 | **263** | 43 | 26.67 |
| **STARlong** | 93.17 | 97.92 | 81.23 | 201 | 6 | 34.88 |

Supplementary Table S28: Benchmark of aligners on real WGS contiguous TGS reads from *Homo sapiens*. Metrics include mapping rate (Map rate, %), minimum alignment identity (Min identity, %), minimum alignment length (Min length), alignment speed (Speed, aligned reads per second), and peak memory usage (RAM, Gb).

| Aligner | Map rate | Min identity | Min length | Speed | RAM |
|---|---|---|---|---|---|
| **VAT** | **100.00** | 62.89 | 44 | **63** | 21.45 |
| **BLASR** | **100.00** | 61.23 | 40 | 1 | 30.33 |
| **GMAP** | 86.95 | 60.77 | **45** | 4 | **14.20** |
| **GraphMap2** | 20.97 | **62.96** | 41 | 2 | 84.37 |
| **ngmlr** | 83.47 | 61.13 | 44 | 13 | 23.78 |
| **Minimap2** | **100.00** | 60.87 | 44 | 30 | 24.16 |
| **STARlong** | 84.25 | 62.81 | 44 | 5 | 33.41 |

Supplementary Table S29: Benchmark of aligners on real CHIP-seq contiguous TGS read from *Homo sapiens*. Metrics include mapping rate (Map rate, %), minimum alignment identity (Min identity, %), minimum alignment length (Min length), alignment speed (Speed, aligned reads per second), and peak memory usage (RAM, Gb).

| Aligner | Map rate | Min identity | Min length | Speed | RAM |
|---|---|---|---|---|---|
| **VAT** | **100.00** | 66.23 | 42 | **420** | 18.78 |
| **BLASR** | **100.00** | 60.11 | 40 | 14 | 30.33 |
| **GMAP** | 98.98 | **67.89** | 20 | 5 | 12.76 |
| **GraphMap2** | 91.03 | 58.41 | 20 | 11 | 53.77 |
| **ngmlr** | 94.89 | 64.78 | 40 | 363 | **11.89** |
| **Minimap2** | 99.03 | 56.99 | **43** | 259 | 21.32 |
| **STARlong** | 95.51 | 59.23 | 42 | 40 | 33.40 |

Supplementary Table S30: Benchmark of aligners on real ATAC-seq contiguous TGS reads from *Homo sapiens*. Metrics include mapping rate (Map rate, %), minimum alignment identity (Min identity, %), minimum alignment length (Min length), alignment speed (Speed, aligned reads per second), and peak memory usage (RAM, Gb).

| Aligner | Map rate | Min identity | Min length | Speed | RAM |
|---|---|---|---|---|---|
| **VAT** | **100.00** | 60.31 | 65 | **277** | 21.89 |
| **BLASR** | **100.00** | 57.78 | 40 | 12 | 30.06 |
| **GMAP** | 12.88 | **61.02** | 51 | 1 | 11.89 |
| **GraphMap2** | 10.92 | 60.10 | 50 | 1 | 78.04 |
| **ngmlr** | 96.21 | 60.78 | 51 | 96 | **10.41** |
| **Minimap2** | **100.00** | 56.04 | **66** | 196 | 19.90 |
| **STARlong** | 77.58 | 59.07 | 56 | 7 | 38.44 |

Supplementary Table S31: Benchmark of aligners on microbiome contiguous TGS reads. Metrics include mapping rate (Map rate, %), minimum alignment identity (Min identity, %), minimum alignment length (Min length), alignment speed (Speed, aligned reads per second), and peak memory usage (RAM, Gb).

| Aligner | Map rate | Min identity | Min length | Speed | RAM |
|---|---|---|---|---|---|
| **VAT** | **100.00** | 65.03 | 43 | **2355** | 19.78 |
| **BLASR** | **100.00** | 62.09 | 39 | 149 | 19.45 |
| **GMAP** | 94.94 | **67.89** | 40 | 13 | 12.03 |
| **GraphMap2** | 0.00 | 0.00 | 0 | 0 | **11.21** |
| **ngmlr** | 63.39 | 59.78 | 39 | 127 | 13.85 |
| **Minimap2** | 99.62 | 61.94 | **44** | 58 | 25.06 |
| **STARlong** | 36.96 | 60.88 | 36 | 7 | 26.67 |

Supplementary Table S32: Benchmark of aligners on contiguous TGS reads from *Caenorhabditis elegans.* Metrics include mapping rate (Map rate, %), minimum alignment identity (Min identity, %), minimum alignment length (Min Iength), alignment speed (Speed, aligned reads per second), and peak memory usage (RAM, Gb).

| Aligner | Map rate | Min identity | Min length | Speed | RAM |
|---|---|---|---|---|---|
| **VAT** | **100.00** | 62.89 | 318 | **893** | 17.23 |
| **BLASR** | **100.00** | 59.04 | 39 | 69 | 27.89 |
| **GMAP** | 99.35 | 59.14 | 71 | 15 | **11.89** |
| **GraphMap2** | 14.01 | 59.08 | 69 | 29 | 66.89 |
| **ngmlr** | 99.13 | 62.77 | 36 | 256 | 13.31 |
| **Minimap2** | 99.80 | **65.89** | 40 | 693 | 23.88 |
| **STARlong** | 95.26 | 60.05 | **320** | 227 | 26.37 |

Supplementary Table S33: Benchmark of aligners on contiguous TGS reads from *Drosophila melanogaster*. Metrics include mapping rate (Map rate, %), minimum alignment identity (Min identity, %), minimum alignment length (Min length), alignment speed (Speed, aligned reads per second), and peak memory usage (RAM, Gb).

| Aligner | Map rate | Min identity | Min length | Speed | RAM |
|---|---|---|---|---|---|
| **VAT** | **100.00** | 65.77 | 50 | **426** | 22.24 |
| **BLASR** | **100.00** | 59.45 | 40 | 26 | 26.99 |
| **GMAP** | 98.03 | **68.04** | 21 | 7 | 13.77 |
| **GraphMap2** | 12.97 | 63.10 | 40 | 19 | 59.14 |
| **ngmlr** | 87.84 | 64.00 | **51** | 54 | **11.04** |
| **Minimap2** | 98.89 | 66.88 | 43 | 412 | 22.84 |
| **STARlong** | 41.19 | 65.32 | 41 | 18 | 26.19 |

Supplementary Table S34: Benchmark of aligners on contiguous TGS reads from *Mus musculus*. Metrics include mapping rate (Map rate, %), minimum alignment identity (Min identity, %), minimum alignment length (Min length), alignment speed (Speed, aligned reads per second), and peak memory usage (RAM, Gb).

| Aligner | Map rate | Min identity | Min length | Speed | RAM |
|---|---|---|---|---|---|
| **VAT** | **100.00** | 59.41 | 62 | **490** | 21.94 |
| **BLASR** | **100.00** | 58.33 | 40 | 14 | 27.31 |
| **GMAP** | 90.77 | 55.76 | 52 | 36 | **12.87** |
| **GraphMap2** | 41.89 | **60.90** | 43 | 8 | 85.43 |
| **ngmlr** | 88.98 | 50.40 | 52 | 41 | **12.87** |
| **Minimap2** | 92.25 | 55.23 | 40 | 128 | 24.10 |
| **STARlong** | 46.28 | 60.11 | **63** | 4 | 48.51 |

Supplementary Table S35: Benchmark of aligners on contiguous TGS reads from *Oryza sativa*. Metrics include mapping rate (Map rate, %), minimum alignment identity (Min identity, %), minimum alignment length (Min length), alignment speed (Speed, aligned reads per second), and peak memory usage (RAM, Gb).

| Aligner | Map rate | Min identity | Min length | Speed | RAM |
|---|---|---|---|---|---|
| **VAT** | **100.00** | 64.04 | **48** | **141** | 17.94 |
| **BLASR** | **100.00** | 60.20 | 46 | 28 | 24.99 |
| **GMAP** | 87.93 | **68.33** | 35 | 8 | **11.30** |
| **GraphMap2** | 60.94 | 61.23 | 44 | 42 | 65.04 |
| **ngmlr** | 94.91 | 62.18 | 44 | 58 | 11.34 |
| **Minimap2** | 99.98 | 60.78 | 45 | 139 | 20.88 |
| **STARlong** | 67.62 | 61.04 | 45 | 43 | 29.11 |

Supplementary Table S36: Benchmark of aligners on contiguous TGS reads from *Arabidopsis thaliana*. Metrics include mapping rate (Map rate, %), minimum alignment identity (Min identity, %), minimum alignment length (Min length), alignment speed (Speed, aligned reads per second), and peak memory usage (RAM, Gb).

| Aligner | Map rate | Min identity | Min length | Speed | RAM |
|---|---|---|---|---|---|
| **VAT** | **100.00** | 64.88 | **42** | **7692** | 16.23 |
| **BLASR** | **100.00** | 61.23 | 26 | 452 | 23.59 |
| **GMAP** | 88.97 | 63.78 | **42** | 185 | **9.98** |
| **GraphMap2** | 89.95 | 60.00 | 31 | 224 | 61.87 |
| **ngmlr** | **100.00** | 63.50 | 22 | 1351 | 12.43 |
| **Minimap2** | 94.87 | 62.00 | 40 | 5929 | 19.03 |
| **STARlong** | 51.04 | **69.87** | 18 | 132 | 29.50 |

Supplementary Table S37: Summary of the split third-generation sequencing (TGS) datasets and simulation commands used in this study. The columns represent the dataset label (Label), sequence continuity status (SCS), data type, source organism (Organism), number of reads (# Reads), average read (Len.), and accession numbers. *H. sapiens* refers to *Homo sapiens*, Gut MG refers to Gut Metagenome, *C. elegans* (*Caenorhabditis elegans*), *D. melan* (*Drosophila melanogaster*), *M. mus* (*Mus musculus*), *O. sativa* (*Oryza sativa*), and *A. thalia* (*Arabidopsis thaliana*). Specifically, TGS-Sim-DS1 is a simulated whole-genome sequencing dataset with an error rate of 12%; TGS-Sim-DS2 is a simulated RNA-Seq dataset with an introduced error rate of 15%. TGS-Sim-DS3 is a simulated Hi-C dataset with an error rate of 15%.

| Label | SCS | Data type | Organism | # Reads | Len. | Accession | Simulation parameters |
|---|---|---|---|---|---|---|---|
| **TGS-Sim-DS2 (PBSIM)** | Split | Simulation | *H. sapiens* | 0.41M | 7800 | N/A | pbsim --data-type CLR --depth 40 --length-mean 3080 --length-sd 2211 --length-min 50 --length-max 50000 --accuracy-mean 0.95 --accuracy-sd 0.11 --accuracy-min 0.7 --difference-ratio 47:38:15 transcriptome_hg38.fa |
| **TGS-DS10** | Split | RNA-Seq | *H. sapiens* | 0.43M | 928 | SRR32923630 | N/A |
| **TGS-DS11** | Split | RNA-Seq | *C. elegans* | 1.1M | 902 | SRR29522715 | N/A |
| **TGS-DS12** | Split | RNA-Seq | *D. melan* | 2.4M | 1113 | SRR32701026 | N/A |
| **TGS-DS13** | Split | RNA-Seq | *M. mus* | 530.9K | 1024 | SRR32517545 | N/A |
| **TGS-DS14** | Split | RNA-Seq | *O. sativa* | 6.5M | 814 | SRR30002035 | N/A |
| **TGS-DS15** | Split | RNA-Seq | *A. thalia* | 42.8K | 513 | ERR14185199 | N/A |

Supplementary Table S38: Summary of long-read split aligners, their software versions, and the specific command-line parameters used for benchmarking in this study.

| Aligner | Version | Command |
|---|---|---|
| **BLASR** | 2012 | -nproc 16 -sam -bestn 10 |
| **GMAP** | 6/24/2024 | -f samse -t 16 |
| **GraphMap2** | 0.6.5 | -t 16 |
| **Minimap2** | 2.30 (r1287) | -ax splice -uf -k14 -t 16 |
| **ngmlr** | 0.2.7 | -x ont -t 16 |
| **STARlong** | 2.7.11b | --runThreadN 16 \<br>--seedSearchStartLmax 14 \<br>--seedPerReadNmax 100000 \<br>--seedPerWindowNmax 100 \<br>--winAnchorMultimapNmax 200 \<br>--outFilterMultimapNmax 100000 \<br>--outFilterMismatchNmax 100000 \<br>--outFilterScoreMin 0 --outFilterScoreMinOverLread 0 \<br>--outFilterMatchNmin 0 --outFilterMatchNminOverLread 0 \<br>--alignIntronMin 20 --alignIntronMax 1000000 \<br>--alignSJoverhangMin 8 --alignSJDBoverhangMin 1 \<br>--alignMatesGapMax 1000000 \<br>--clip3pAfterAdapterNbases 1 --clip5pNbases 0 \<br>--chimOutType WithinBAM SoftClip \<br>--outSAMtype SAM \<br>--limitBAMsortRAM 12000000000 |
| **VAT** | 0.0.1 | -p 16 -splice --long |

Supplementary Table S39: Benchmark of long-read aligners on simulated split TGS reads from *Homo sapiens*. Metrics include mapping rate (Map rate, %), proportions of contiguous alignments (Contiguous, %), proportions of split alignments (Split, %), accuracy (%), proportion of known split events (Known, %), alignment speed (Speed, aligned reads per second), and peak memory usage (RAM, Gb).

| Aligner | Map rate | Contiguous | Split | Accuracy | Known | Speed | RAM |
|---|---|---|---|---|---|---|---|
| **VAT** | **98.02** | 22.91 | **75.11** | **86.80** | **83.36** | **964** | 24.41 |
| **BLASR** | 94.49 | 27.89 | 66.60 | 78.82 | 71.45 | 79 | 22.59 |
| **GMAP** | 74.77 | 17.76 | 57.01 | 81.29 | 76.83 | 23 | **10.04** |
| **GraphMap2** | 94.50 | 20.30 | 74.20 | 76.56 | 71.74 | 19 | 101.11 |
| **ngmlr** | 54.80 | 22.48 | 32.32 | 53.00 | 20.80 | 627 | 11.88 |
| **Minimap2** | 94.61 | 21.70 | 72.91 | 85.95 | 82.33 | 659 | 25.78 |
| **STARlong** | 85.06 | **35.15** | 49.91 | 84.70 | 74.95 | 155 | 22.79 |

Supplementary Table S40: Benchmark of long-read aligners on real TGS RNA-seq reads from *Homo sapiens*. Metrics include mapping rate (Map rate, %), proportions of contiguous alignments (Contiguous, %), proportions of split alignments (Split, %), proportion of known split events (Known, %), alignment speed (Speed, aligned reads per second), and peak memory usage (RAM, Gb).

| Aligner | Map rate | Contiguous | Split | Known | Speed | RAM |
|---|---|---|---|---|---|---|
| **VAT** | **96.81** | 25.97 | **70.84** | **84.46** | **899** | 23.33 |
| **BLASR** | 95.68 | 34.96 | 60.72 | 73.42 | 86 | 100.89 |
| **GMAP** | 90.69 | 40.88 | 49.81 | 73.22 | 84 | **9.11** |
| **GraphMap2** | 59.98 | 30.11 | 29.87 | 66.82 | 3 | 64.87 |
| **ngmlr** | 54.97 | **49.89** | 5.08 | 40.16 | 14 | 12.89 |
| **Minimap2** | 95.48 | 35.03 | 60.45 | 77.70 | 789 | 24.91 |
| **STARlong** | 87.25 | 40.89 | 46.36 | 81.77 | 53 | 38.14 |

Supplementary Table S41: Benchmark of long-read aligners on TGS RNA-seq reads from *Caenorhabditis elegans*. Metrics include mapping rate (Map rate, %), proportions of contiguous alignments (Contiguous, %), proportions of split alignments (Split, %), proportion of known split events (Known, %), alignment speed (Speed, aligned reads per second), and peak memory usage (RAM, Gb).

| Aligner | Map rate | Contiguous | Split | Known | Speed | RAM |
|---|---|---|---|---|---|---|
| **VAT** | **100.00** | 11.03 | **88.97** | 88.80 | **16129** | 20.89 |
| **BLASR** | **100.00** | **65.00** | 35.00 | 53.68 | 810 | 23.14 |
| **GMAP** | 87.99 | 8.49 | 79.50 | 77.78 | 356 | **9.89** |
| **GraphMap2** | 86.20 | 13.62 | 72.58 | **91.73** | 397 | 100.91 |
| **ngmlr** | 58.09 | 48.03 | 10.06 | 20.69 | 1208 | 12.33 |
| **Minimap2** | 80.98 | 9.09 | 71.89 | 85.65 | 10176 | 25.78 |
| **STARlong** | 67.28 | 13.72 | 53.56 | 83.08 | 18 | 24.53 |

Supplementary Table S42: Benchmark of long-read aligners on TGS RNA-seq reads from *Drosophila melanogaster*. Metrics include mapping rate (Map rate, %), proportions of contiguous alignments (Contiguous, %), proportions of split alignments (Split, %), proportion of known split events (Known, %), alignment speed (Speed, aligned reads per second), and peak memory usage (RAM, Gb).

| Aligner | Map rate | Contiguous | Split | Known | Speed | RAM |
|---|---|---|---|---|---|---|
| **VAT** | **99.91** | 41.03 | **58.88** | **88.14** | **47610** | 22.23 |
| **BLASR** | 98.96 | 61.92 | 37.04 | 67.43 | 3690 | 28.97 |
| **GMAP** | 84.19 | 36.94 | 47.25 | 85.44 | 76 | 11.43 |
| **GraphMap2** | 90.89 | 41.87 | 49.02 | 83.51 | 2390 | 65.13 |
| **ngmlr** | 85.02 | **63.31** | 21.71 | 37.45 | 443 | **10.96** |
| **Minimap2** | 91.71 | 38.60 | 53.11 | 85.14 | 29568 | 25.30 |
| **STARlong** | 73.21 | 29.34 | 43.86 | 84.10 | 158 | 27.41 |

Supplementary Table S43: Benchmark of long-read aligners on TGS RNA-seq reads from *Mus musculus*. Metrics include mapping rate (Map rate, %), proportions of contiguous alignments (Contiguous, %), proportions of split alignments (Split, %), proportion of known split events (Known, %), alignment speed (Speed, aligned reads per second), and peak memory usage (RAM, Gb).

| Aligner | Map rate | Contiguous | Split | Known | Speed | RAM |
|---|---|---|---|---|---|---|
| **VAT** | **100.00** | 20.03 | **79.97** | **83.81** | **17241** | 23.33 |
| **BLASR** | **100.00** | 28.98 | 71.02 | 78.12 | 490 | 26.78 |
| **GMAP** | 94.96 | 23.96 | 71.00 | 67.26 | 238 | 11.89 |
| **GraphMap2** | 94.98 | 25.01 | 69.97 | 83.59 | 303 | 65.13 |
| **ngmlr** | 97.59 | **62.69** | 34.90 | 48.31 | 4866 | **10.78** |
| **Minimap2** | 99.81 | 23.03 | 76.78 | 83.07 | 6836 | 24.98 |
| **STARlong** | 80.04 | 35.39 | 44.64 | 77.26 | 292 | 37.06 |

Supplementary Table S44: Benchmark of long-read aligners on TGS RNA-seq reads from *Oryza sativa*. Metrics include mapping rate (Map rate, %), proportions of contiguous alignments (Contiguous, %), proportions of split alignments (Split, %), proportion of known split events (Known, %), alignment speed (Speed, aligned reads per second), and peak memory usage (RAM, Gb).

| Aligner | Map rate | Contiguous | Split | Known | Speed | RAM |
|---|---|---|---|---|---|---|
| **VAT** | **100.00** | 54.89 | 45.11 | **80.04** | **52632** | 23.03 |
| **BLASR** | **100.00** | 47.98 | **52.02** | 61.58 | 2618 | 28.33 |
| **GMAP** | 93.01 | 50.04 | 42.97 | 78.85 | 911 | **10.91** |
| **GraphMap2** | 98.58 | **74.87** | 23.71 | 54.44 | 1641 | 62.77 |
| **ngmlr** | 26.01 | 23.03 | 2.98 | 43.63 | 8387 | 11.04 |
| **Minimap2** | 99.47 | 71.97 | 27.50 | 67.15 | 17454 | 22.95 |
| **STARlong** | 93.00 | 43.46 | 49.54 | 77.23 | 141 | 27.11 |

Supplementary Table S45: Benchmark of long-read aligners on TGS RNA-seq reads from *Arabidopsis thaliana*. Metrics include mapping rate (Map rate, %), proportions of contiguous alignments (Contiguous, %), proportions of split alignments (Split, %), proportion of known split events (Known, %), alignment speed (Speed, aligned reads per second), and peak memory usage (RAM, Gb).

| Aligner | Map rate | Contiguous | Split | Known | Speed | RAM |
|---|---|---|---|---|---|---|
| **VAT** | **100.00** | 48.03 | **51.97** | **86.41** | **125000** | 21.06 |
| **BLASR** | 92.80 | 46.89 | 45.91 | 47.67 | 11364 | 28.20 |
| **GMAP** | 89.98 | 49.03 | 40.95 | 70.11 | 4093 | 12.78 |
| **GraphMap2** | 93.88 | 42.97 | 50.91 | 72.03 | 6345 | 61.50 |
| **ngmlr** | 59.39 | 46.01 | 13.38 | 55.38 | 13395 | **9.95** |
| **Minimap2** | 90.49 | 39.88 | 50.61 | 77.29 | 69615 | 23.20 |
| **STARlong** | 99.52 | **89.32** | 10.20 | 77.98 | 29 | 36.45 |

Supplementary Table S46: Summary of protein homology search tools, their software versions, and the specific command-line parameters used for benchmarking in this study.

| Aligner | Version | Command |
|---|---|---|
| **BLAST** | 2.16.0 | blastp<br>-outfmt 6 -num_threads 8 -max_target_seqs 5 -evalue |
| **DIAMOND** | 2.1.12 | -p 8 –fast/ultra-sensitive -k 5 |
| **MMseqs2** | 17-b804f | tmp --threads 8 -e --max-seqs 5 -s 1/7.5 |
| **RAPSearch2** | 2.22 | -z 4 -m 8 -b 5 |
| **VAT** | 0.0.1 | protein -p 8 -e -k 5 –fast/sensitive |

Supplementary Table S47: Summary of DNA homology search tools, their software versions, and the specific command-line parameters used for benchmarking in this study.

| Aligner | Version | Command |
|---|---|---|
| **BLAST** | 2.16.0 | blastn<br>-outfmt 6 -num_threads 16 -max_target_seqs 2 |
| **pblat** | 2.1.12 | -threads=16 -minIdentity=90 -minScore=30 -tileSize=8 -stepSize=5 -out=blast8 |
| **VAT** | 0.0.1 | dna -p 16 -e -k 2 –dnah |

Supplementary Table S48: Benchmark of protein homology search tools across varying E-value thresholds. Metrics include precision, sensitivity, F1 score (F1), alignment speed (Speed, aligned queries per second), and peak memory usage (RAM, Gb).

| Aligner | E-value | Precision | Sensitivity | F1 | Speed | RAM |
|---|---|---|---|---|---|---|
| **BLASTP** | 1 | 0.756 | **0.730** | 0.743 | 47 | 0.161 |
| **BLASTP** | 0.1 | 0.922 | 0.679 | **0.782** | 48 | 0.164 |
| **BLASTP** | 0.01 | 0.960 | 0.640 | 0.768 | 50 | 0.161 |
| **BLASTP** | 0.001 | 0.970 | 0.603 | 0.744 | 51 | 0.162 |
| **BLASTP** | 0.0001 | 0.973 | 0.566 | 0.716 | 54 | 0.161 |
| **BLASTP** | 1.00E-05 | 0.977 | 0.532 | 0.689 | 54 | 0.158 |
| **BLASTP** | 1.00E-06 | 0.979 | 0.500 | 0.662 | 54 | 0.153 |
| **BLASTP** | 1.00E-07 | 0.980 | 0.472 | 0.637 | 54 | **0.144** |
| **VAT sensitive** | 1 | 0.873 | 0.679 | 0.764 | 341 | 0.558 |
| **VAT sensitive** | 0.1 | 0.932 | 0.657 | 0.773 | 348 | 0.549 |
| **VAT sensitive** | 0.01 | 0.958 | 0.636 | 0.765 | 347 | 0.549 |
| **VAT sensitive** | 0.001 | 0.967 | 0.602 | 0.742 | 347 | 0.553 |
| **VAT sensitive** | 0.0001 | 0.971 | 0.578 | 0.729 | 346 | 0.539 |
| **VAT sensitive** | 1.00E-05 | 0.977 | 0.543 | 0.699 | 345 | 0.538 |
| **VAT sensitive** | 1.00E-06 | 0.978 | 0.512 | 0.671 | 344 | 0.537 |
| **VAT sensitive** | 1.00E-07 | 0.981 | 0.471 | 0.637 | 344 | 0.539 |
| **MMseqs2 s7.5** | 1 | 0.880 | 0.661 | 0.755 | 223 | 1.015 |
| **MMseqs2 s7.5** | 0.1 | 0.941 | 0.640 | 0.762 | 239 | 1.013 |
| **MMseqs2 s7.5** | 0.01 | 0.960 | 0.622 | 0.755 | 241 | 1.011 |
| **MMseqs2 s7.5** | 0.001 | 0.972 | 0.591 | 0.735 | 243 | 1.011 |
| **MMseqs2 s7.5** | 0.0001 | 0.973 | 0.550 | 0.703 | 246 | 1.012 |
| **MMseqs2 s7.5** | 1.00E-05 | 0.974 | 0.522 | 0.680 | 248 | 1.01 |
| **MMseqs2 s7.5** | 1.00E-06 | 0.977 | 0.490 | 0.653 | 248 | 1.002 |
| **MMseqs2 s7.5** | 1.00E-07 | 0.980 | 0.461 | 0.627 | 249 | 1.005 |
| **DIAMOND ultra-sensitive** | 1 | 0.888 | 0.663 | 0.759 | 301 | 0.321 |
| **DIAMOND ultra-sensitive** | 0.1 | 0.940 | 0.649 | 0.768 | 319 | 0.312 |
| **DIAMOND ultra-sensitive** | 0.01 | 0.961 | 0.626 | 0.758 | 324 | 0.331 |
| **DIAMOND ultra-sensitive** | 0.001 | 0.971 | 0.592 | 0.736 | 343 | 0.311 |
| **DIAMOND ultra-sensitive** | 0.0001 | 0.975 | 0.559 | 0.711 | 343 | 0.308 |
| **DIAMOND ultra-sensitive** | 1.00E-05 | 0.977 | 0.527 | 0.685 | 344 | 0.306 |
| **DIAMOND ultra-sensitive** | 1.00E-06 | 0.979 | 0.497 | 0.659 | 344 | 0.31 |
| **DIAMOND ultra-sensitive** | 1.00E-07 | 0.981 | 0.469 | 0.635 | 347 | 0.305 |
| **VAT fast** | 1 | 0.920 | 0.370 | 0.528 | 3301 | 0.625 |
| **VAT fast** | 0.1 | 0.944 | 0.366 | 0.527 | 3322 | 0.626 |
| **VAT fast** | 0.01 | 0.955 | 0.361 | 0.524 | 3330 | 0.611 |
| **VAT fast** | 0.001 | 0.963 | 0.354 | 0.518 | 3335 | 0.611 |
| **VAT fast** | 0.0001 | 0.968 | 0.345 | 0.509 | 3341 | 0.604 |
| **VAT fast** | 1.00E-05 | 0.971 | 0.336 | 0.499 | 3341 | 0.613 |

Continue to next page

| | | | | | | |
|---|---|---|---|---|---|---|
| **VAT fast** | 1.00E-06 | 0.972 | 0.327 | 0.489 | 3352 | 0.615 |
| **VAT fast** | 1.00E-07 | 0.975 | 0.318 | 0.480 | 3352 | 0.609 |
| **MMseqs2 s1** | 1 | 0.980 | 0.219 | 0.358 | 2488 | 0.91 |
| **MMseqs2 s1** | 0.1 | 0.981 | 0.218 | 0.357 | 2491 | 0.903 |
| **MMseqs2 s1** | 0.01 | 0.981 | 0.218 | 0.357 | 2491 | 0.905 |
| **MMseqs2 s1** | 0.001 | 0.982 | 0.217 | 0.355 | 2493 | 0.901 |
| **MMseqs2 s1** | 0.0001 | 0.983 | 0.215 | 0.353 | 2503 | 0.898 |
| **MMseqs2 s1** | 1.00E-05 | 0.983 | 0.213 | 0.350 | 2506 | 0.902 |
| **MMseqs2 s1** | 1.00E-06 | 0.984 | 0.211 | 0.347 | 2510 | 0.899 |
| **MMseqs2 s1** | 1.00E-07 | 0.984 | 0.208 | 0.343 | 2512 | 0.905 |
| **DIAMOND fast** | 1 | 0.982 | 0.187 | 0.314 | 3301 | 0.161 |
| **DIAMOND fast** | 0.1 | 0.983 | 0.186 | 0.313 | 3305 | 0.162 |
| **DIAMOND fast** | 0.01 | 0.983 | 0.186 | 0.313 | 3317 | 0.166 |
| **DIAMOND fast** | 0.001 | 0.984 | 0.185 | 0.311 | 3325 | 0.171 |
| **DIAMOND fast** | 0.0001 | 0.984 | 0.184 | 0.310 | 3335 | 0.161 |
| **DIAMOND fast** | 1.00E-05 | 0.984 | 0.183 | 0.309 | 3346 | 0.158 |
| **DIAMOND fast** | 1.00E-06 | **0.985** | 0.181 | 0.306 | 3356 | 0.159 |
| **DIAMOND fast** | 1.00E-07 | **0.985** | 0.180 | 0.304 | **3361** | 0.157 |
| **RAPSearch2** | 1 | 0.813 | 0.482 | 0.605 | 1231 | 0.836 |
| **RAPSearch2** | 0.1 | 0.928 | 0.472 | 0.626 | 1246 | 0.833 |
| **RAPSearch2** | 0.01 | 0.931 | 0.471 | 0.626 | 1249 | 0.814 |
| **RAPSearch2** | 0.001 | 0.931 | 0.471 | 0.626 | 1250 | 0.811 |
| **RAPSearch2** | 0.0001 | 0.931 | 0.471 | 0.626 | 1256 | 0.809 |
| **RAPSearch2** | 1.00E-05 | 0.931 | 0.471 | 0.626 | 1271 | 0.822 |
| **RAPSearch2** | 1.00E-06 | 0.931 | 0.471 | 0.626 | 1271 | 0.811 |
| **RAPSearch2** | 1.00E-07 | 0.931 | 0.471 | 0.626 | 1278 | 0.801 |

Supplementary Table S49: Benchmark of DNA homology search tools across varying E-value thresholds. Metrics include precision, sensitivity, F1 score (F1), alignment speed (Speed, aligned queries per second), and peak memory usage (Memory, Gb).

| Aligner | E-value | Precision | Sensitivity | F1 | Speed | RAM |
|---|---|---|---|---|---|---|
| **BLASTN** | 1 | 0.540 | 0.310 | 0.394 | 13 | 0.471 |
| **BLASTN** | 0.1 | 0.545 | 0.303 | 0.389 | 13 | 0.431 |
| **BLASTN** | 0.01 | 0.550 | 0.300 | 0.388 | 14 | 0.445 |
| **BLASTN** | 0.001 | 0.554 | 0.298 | 0.388 | 15 | 0.433 |
| **BLASTN** | 0.0001 | 0.558 | 0.296 | 0.387 | 14 | 0.436 |
| **BLASTN** | 1.00E-05 | 0.562 | 0.294 | 0.386 | 15 | **0.422** |
| **BLASTN** | 1.00E-06 | **0.567** | 0.289 | 0.383 | 15 | 0.423 |
| **VAT** | 1 | 0.530 | **0.350** | **0.422** | 27 | 1.461 |
| **VAT** | 0.1 | 0.540 | 0.320 | 0.402 | 28 | 1.423 |
| **VAT** | 0.01 | 0.548 | 0.310 | 0.396 | 28 | 1.433 |
| **VAT** | 0.001 | 0.553 | 0.304 | 0.392 | 29 | 1.413 |
| **VAT** | 0.0001 | 0.556 | 0.300 | 0.390 | 29 | 1.411 |
| **VAT** | 1.00E-05 | 0.560 | 0.297 | 0.388 | **30** | 1.423 |
| **VAT** | 1.00E-06 | 0.564 | 0.294 | 0.387 | 29 | 1.426 |
| **pblat** | 1 | 0.539 | 0.304 | 0.389 | 10 | 0.994 |
| **pblat** | 0.1 | 0.540 | 0.303 | 0.388 | 10 | 0.991 |
| **pblat** | 0.01 | 0.542 | 0.302 | 0.388 | 10 | 0.992 |
| **pblat** | 0.001 | 0.544 | 0.301 | 0.388 | 10 | 0.976 |
| **pblat** | 0.0001 | 0.548 | 0.300 | 0.388 | 10 | 0.983 |
| **pblat** | 1.00E-05 | 0.550 | 0.298 | 0.387 | 10 | 0.977 |
| **pblat** | 1.00E-06 | 0.551 | 0.297 | 0.386 | 10 | 0.992 |

Supplementary Table S50: Comparison of VAT-fast, DIAMOND-fast, and MMseqs2-s1 in aligning two soil metagenomic datasets. Metrics include alignment rate (%), aligned matches, queries aligned, alignment speed (Speed, aligned queries per second), and peak memory usage (RAM, Gb).

| Datasets | Aligner | Alignment rate | Aligned matches | Aligned queries | Speed | RAM |
|---|---|---|---|---|---|---|
| **ERR1872095** | VAT | **99.28** | **1898994** | **65940** | 13 | 28.77 |
| | DIAMOND | 96.99 | 1641135 | 64426 | **14** | **10.71** |
| | MMseqs2 | 98.51 | 1652341 | 65429 | 9 | 324.65 |
| **ERR1873275** | VAT | **99.36** | **2528103** | **90930** | 21 | 27.81 |
| | DIAMOND | 98.05 | 2344125 | 89741 | **23** | **10.79** |
| | MMseqs2 | 98.85 | 2319446 | 90463 | 13 | 351.73 |

Supplementary Table S51: Summary of reference genome assemblies used for whole-genome alignment, including version, total genome size, and number of reference sequences.

| Organism | Version | Size | # References |
|---|---|---|---|
| ***Homo sapiens*** | GRCh38.p7 (GCA_000001405.22) | 3.088 Gb | 525 |
| ***Pan troglodytes*** | Pan_troglodytes-2.1.4 (GCA_000001515.4) | 3.31 Gb | 24128 |
| ***Arabidopsis thaliana*** | TAIR10 (GCF_000001735.3) | 120 Mb | 7 |
| ***Arabidopsis lyrata*** | v.1.0 (GCF_000004255.1) | 207 Mb | 695 |

Supplementary Table S52: Summary of whole genome alignment tools, their software versions, and the specific command-line parameters used for benchmarking in this study.

| Aligner | Version | Command |
|---|---|---|
| **MUMmer4** | 4.0.1 | nucmer -c 100 -t 32 |
| **VAT** | 0.0.1 | -p 32 --wga |

Supplementary Table S53: Comparison of genome alignment coverage between VAT and MUMmer4 across varying sequence similarity levels (%) between *Homo sapiens* and *Pan troglodyte*. Metrics include MUMmer4 coverage (%) and VAT coverage (%).

| Similarity | MUMmer4 | VAT |
|---|---|---|
| **88.00** | 86.32 | 88.47 |
| **90.00** | 86.20 | 87.76 |
| **92.00** | 86.05 | 86.73 |
| **94.00** | 85.75 | 86.09 |
| **96.00** | 84.58 | 85.45 |
| **98.00** | 55.10 | 55.70 |

Supplementary Table S54: Comparison of genome alignment coverage between VAT and MUMmer4 across varying sequence similarity levels (%) between *Arabidopsis thaliana* and *Arabidopsis lyrata*. Metrics include MUMmer4 coverage (%) and VAT coverage (%).

| Similarity | MUMmer4 | VAT |
|---|---|---|
| **70.00** | 52.71 | 54.23 |
| **75.00** | 51.64 | 53.12 |
| **80.00** | 51.59 | 51.87 |
| **85.00** | 35.25 | 34.66 |
| **90.00** | 15.66 | 15.86 |
| **95.00** | 0.46 | 0.62 |

Supplementary Table S55: Whole-genome alignment performance of VAT and MUMmer4 on *Homo sapiens (H. sapiens)* versus *Pan troglodytes (P. troglodytes)* and *Arabidopsis thaliana (A. thaliana)* versus *Arabidopsis lyrata (A. lyrata)*. Metrics include total runtime (Time, minutes) and peak memory usage (RAM, Gb).

| Aligner | ***H. sapiens*** **versus** ***P. troglodytes*** | | ***A. thaliana*** **versus** ***A. lyrata*** | |
|---|---|---|---|---|
| | **Time** | **RAM** | **Time** | **RAM** |
| **VAT** | **43** | 73.76 | **2** | 8.04 |
| **MUMmer4** | 142 | **57.66** | 5 | **2.98** |
